# Ambient Pollution Components and Sources are Associated with Hippocampal Architecture and Memory in Pre-Adolescents

**DOI:** 10.1101/2025.06.05.658105

**Authors:** Michael A. Rosario, Kirthana Sukumaran, Katherine L. Bottenhorn, Alethea de Jesus, Carlos Cadenas-Iniguez, Hedyeh Ahmadi, Rima Habre, Shermaine Abad, Jacob G. Pine, Deanna M. Barch, Joel Schwartz, Daniel A. Hackman, Jiu-Chan Chen, Megan M. Herting

## Abstract

**Background:** Ambient air pollution poses significant risks to brain health. The hippocampus may be particularly vulnerable, yet the extent to which it is impacted in children remains unclear.

**Methods:** Using partial least squares correlation, we cross-sectionally analyzed air pollution, brain, and cognitive data from the Adolescent Brain Cognitive Development Study to examine how multi-pollutant exposure influences hippocampal structure and memory in 9-11-year-olds (n= 7,940). Annual average air pollution exposures included PM_2.5_ (total mass, 15 components, and 6 source factors), NO_2_, and 8-hour maximum O_3_. Hippocampal outcomes included microstructure measured using Restriction Spectrum Imaging and hippocampus longitudinal-axis (i.e., head, body, tail) volumes. We examined hippocampal-dependent list-learning using the Rey Auditory Verbal Learning Test. Models were adjusted for demographic, socioeconomic, and neuroimaging factors.

**Findings:** PM_2.5_ total mass was associated with hippocampal microstructure, but not long-axis volume or list-learning ability. Component and source analyses provided greater specificity: higher bromine, sulfate, and vanadium exposure was related to microstructure (72% shared variance), while higher copper and zinc exposure correlated with smaller left head and right body and tail volumes (75% shared variance). Source models implicated biomass burning and traffic pollution in microstructure (61% and 32% shared variance) and industrial and traffic sources in smaller hippocampal volumes (77% shared variance). Higher exposure to several components were also linked to poorer list-learning (67% shared variance).

**Discussion:** Co-exposure to multiple pollutants is linked to differences in hippocampal structure and memory, showing that associations are driven not only by PM_2.5_ total mass but also by specific components and sources. This evidence underscores the necessity of targeting source-specific (e.g., biomass burning, traffic, and industrial emissions) and constituent components (e.g., metals) of air pollution during critical developmental windows to safeguard brain health.

## Introduction

Exposure to ambient fine particulate matter (PM_2.5_; defined by an aerodynamic diameter of 2.5μm or less) is a pressing challenge to human health, with neurotoxic effects that pose significant risks across the lifespan ^1–4^. PM_2.5_ exposure occurs via inhalation with downstream systemic and neuroinflammatory pathways, and it co-occurs with correlated gases and other pollutants that may index shared sources and exposure contexts ^1,2,15^ Emerging evidence strongly suggests PM_2.5_ exposure threatens neurodevelopment ^5^ and may impair cognitive functioning ^6,7^. Other pollutants, including nitrogen dioxide (NO_2_) and ground-level ozone (8-hour maximum; O_3_)^8^, have also been linked to structural and functional changes in the developing brain ^5^, including cognitive functioning^7,9,10^. While the U.S. Environmental Protection Agency (EPA) regulates these three criteria pollutants under the Clean Air Act ^11^, recent evidence indicates that brain structural and functional differences occur in youth even at exposure levels below current regulatory standards ^5^. Moreover, PM_2.5_ standards focus on quantifying the mass of fine airborne particles but offer no insight into PM_2.5_’s chemical composition or sources ^12,13^. That is, PM_2.5_ is a heterogeneous mixture of organic compounds, metals, nitrates, and sulfates, with the composition substantially varying based on sources, such as biomass burning, traffic, and industrial exposures ^14^. Additionally, several prominent components found in PM_2.5_ can impact the brain through multiple pathways, including direct translocation along the olfactory and trigeminal nerves ^15^, systemic circulation via alveolar-capillary transfer in the lungs ^16^, or activation of hypothalamic-pituitary-adrenal (HPA) axis dysregulation ^17^. Identifying how co-exposure to multiple air pollutants impacts the developing brain is essential for understanding the foundations of cognitive health along the lifespan ^18^ as well as guiding evidence-based air quality policies and protective public health measures.

The hippocampus is a neural structure central to episodic memory and cognitive development ^19–21^. Hippocampal development during childhood supports age-related gains in learning and recall that are foundational for academic and everyday functioning ^19,21^. Its structural development has been extensively characterized across childhood and adolescence ^22–24^, with evidence of functional and structural differentiation along its longitudinal (anterior-posterior) axis ^25–28^. Brain development during this period is rapid and involves ongoing structural and microstructural refinement ^20,22,28^, increasing sensitivity to environmental toxicants ^4,22,24^. Most studies examining the effects of air pollution on the hippocampus during childhood and adolescence have not examined regional volume differences along the hippocampal long-axis, primarily focusing on total hippocampal volume in relation to PM_2.5_ total mass ^29–32^, and rarely its relationship to specific components or sources of PM_2.5_ ^33^. Complementary to this understanding is examining how air pollution influences hippocampal gray matter microstructure, as the hippocampus is composed of distinct cell types with specialized functions ^34^. Diffusion-weighted imaging techniques offer a sensitive means of assessing microstructural properties of brain tissue composition that may reflect differences in cellular density, dendritic complexity, and glial proliferation within the hippocampus ^35,36^. Examining how hippocampal microstructure and which long-axis subregions are impacted in relation to air pollution thus provides an opportunity to detect subtle differences associated with exposure. Importantly, these relationships may differ by pollutant type, reflecting distinct neurotoxic pathways by which air pollution may impact the brain ^17,37^. In sum, assessing both hippocampal microstructure and long-axis subregional structure offers a critical opportunity to detect early signatures of pollution-related neurodevelopmental vulnerability.

Prior animal and human studies show structural and functional effects of air pollution on the hippocampus. Adult male mice and rats exposed to PM_2.5_ or traffic-related air pollution (TRAP) exhibit impaired spatial learning and memory, reduced dendritic complexity, increased microglial activation, disrupted neurogenesis, and compromised blood-brain barrier integrity ^38,39^. Developmental exposure studies similarly report altered microglial and astroglial responsivity ^40,41^, though some studies reported no hippocampal-dependent deficits despite similar exposure windows ^42^. In humans, prenatal exposures to lower NO_2_ and higher PM_10_ was associated with smaller hippocampal volume in infancy ^43^. Prenatal exposure to air pollution is also associated with smaller hippocampal volumes in late childhood ^44,45^ and with delayed hippocampal growth from 4-10-years-old ^30^. Yet, other studies have failed to find such an association in pre-adolescents ^29,31^. These discrepancies may arise from differences in samples, analytic methods, exposure timing, hippocampal assessment, and regional pollution composition ^46^. Consequently, it is essential that multi-pollutant and source-specific models are considered ^47^ to more accurately determine the impact of PM_2.5_ components and their sources on hippocampal architecture and behavior. Given these differences, whether the pollutants that impact hippocampal structure likewise influence learning and memory remains an open question.

The aforementioned studies examining air pollution’s link to brain structure have either not examined ^29,32,33,43–45,48^ or have produced null relationships ^31^ linking air pollution to memory performance. Among highly exposed urban youth, higher levels of pollution have been linked to impaired attention, short-term memory ability, and learning performance ^49^ and a recent meta-analysis showed that greater PM_2.5_ total mass was associated with poorer working memory performance among children ^7^. Additional work has further emphasized the importance of considering the composition and sources of pollution on cognition. Findings from the BREATHE cohort showed that exposure to black carbon and PM_2.5_ was associated with impaired working memory over 12 months ^50^. More recently, we also found that air pollution composition is associated with distinct domains of cognitive functioning in late childhood, with higher annual concentrations of ammonium nitrates being particularly harmful for memory performance ^10^. Taken together, while we have begun to understand the impact of air pollution on gross hippocampal structure and other cognitive abilities, our understanding of its link to hippocampal-dependent memory is lacking.

In this study, we assessed associations between hippocampal structure and annual averages of PM_2.5_, NO_2_ and 8-hour maximum ground-level O_3_, in a large cohort of 9-10-year-olds from the U.S. Adolescent Brain Cognitive Development^SM^ (ABCD) Study ^51,52^. To better characterize PM_2.5_ effects, we examined total mass, constituent components, and emission sources ^10,53^, as these represent distinct dimensions of exposure that may differentially relate to brain and cognitive development (Figure 1 A-D). We assessed hippocampal architecture using complementary MRI measures of microstructure ^55–58^ and long-axis volume ^59–61^, which capture distinct aspects of tissue organization ^55,57,58^ relevant to memory function. Given the highly interrelated nature of air pollution exposures and brain measures, we employed partial least squares correlation to identify multivariate covariance patterns linking environmental exposures, hippocampal structure, and episodic memory performance. We hypothesized that higher air pollution exposure would be associated with altered hippocampal microstructure and smaller long-axis volumes, and to poorer memory performance.

**Figure 1.**
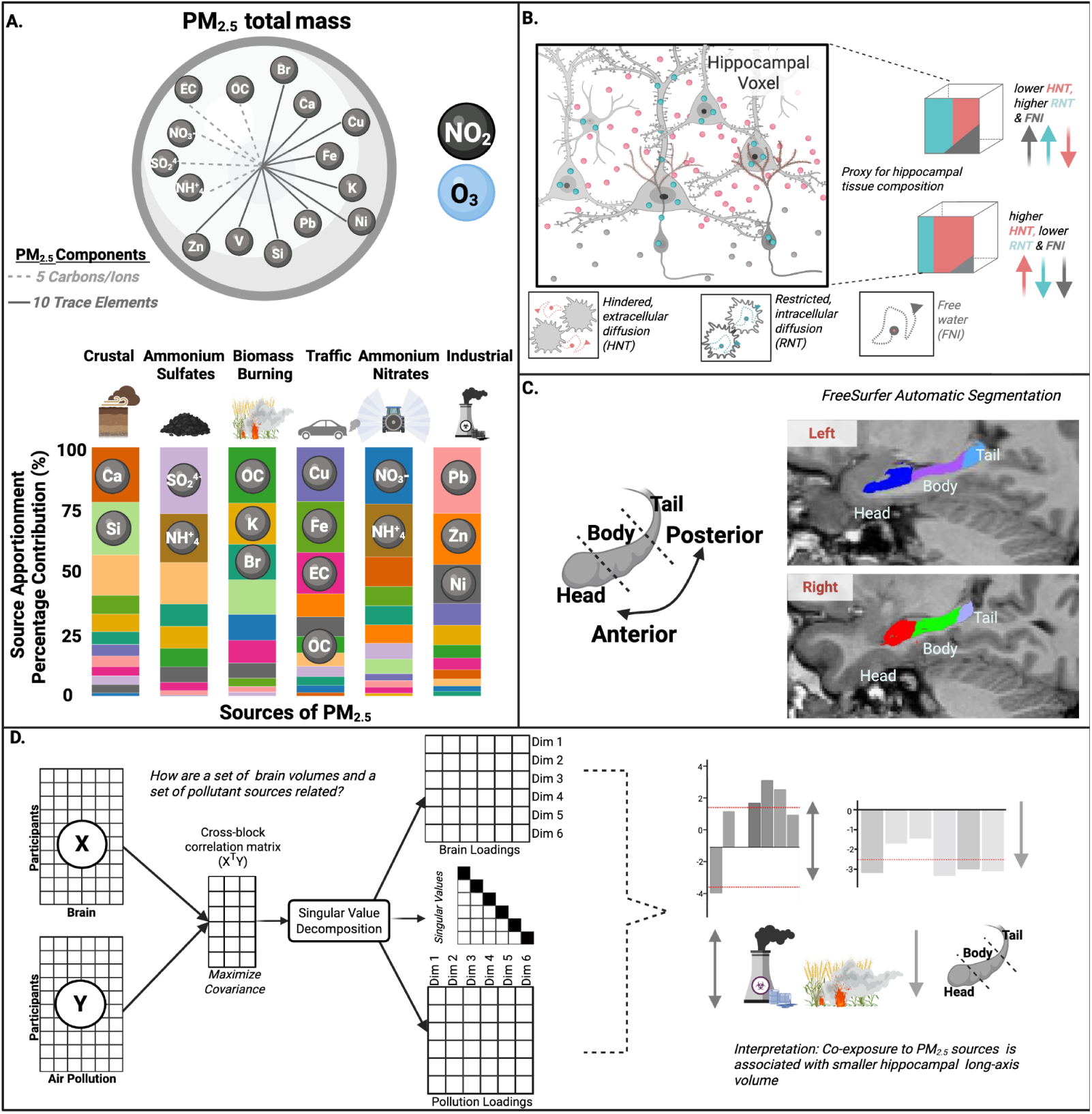
Schematic of exposures and outcomes for air pollution exposures and hippocampal analyses in 9-11-year-olds. **A,** Annual average PM_2.5_ total mass, NO_2_ and O_3_ (8-hour max), fifteen PM_2.5_ components and six PM_2.5_-related source factors derived from individual ABCD cohort participant-level PM_2.5_ component estimates using Positive Matrix Factorization^10^. Source-factors represent a mixture of component exposures: crustal materials driven by Si and Ca; ammonium sulfates driven by NH_4_^+^, SO_4_^2-^; biomass burning driven by OC, K, and Br; traffic emissions driven by EC, Fe, Cu; ammonium nitrates driven by NH_4_^+^ and NO_3_^-^; and industrial/residual fuel burning driven by Pb, Zn, and Ni. **B.** Hippocampal gray matter microstructure as measured by restriction spectrum imaging (RSl), including voxel-level estimates of total hindered, extracellular diffusion (HNT), total restricted, intracellular diffusion (RNT), and free waler (FNI) normalized signal fractions. Compromised microstructure may include: lower external hindered diffusion indicating a less tortuous, more open extracellular space with fewer cellular obstacles, while higher free water diffusion may suggest increased extracellular fluid in expanded spaces. **C.** Hippocampal volumes for head, body and tail along the hippocampal long-axis (anterior-to-posterior) using FreeSurfer. **D.** Partial Least Square Correlation (PLSC) analyses overview, where the cross-block correlation matrix is decomposed via singular value decomposition (SVO) to extract latent dimensions that maximize shared variance between X (e.g., air pollution exposures) and Y (e.g. hippocampal structure). Abbreviations: Br: bromine, Ca: calcium, Cu: copper, EC: elemental carbon, Fe: iron, K: potassium, NH^+^_4_: ammonium, Ni: nickel, NO_2_: nitrogen dioxide, NO_3_^-^: nitrate, O_3_: ozone, OC: organic carbon, Pb: lead, PM_2.5_ mass: fine particulate matter, PMF: positive matrix factorization, Si: silicon, SO_4_^2-^: sulfate, V: vanadium. Zn: zinc, RSI: restriction spectrum imaging; LRNT: left restricted normalized total fraction, RRNT: right restricted normalized total fraction, LHNT: left hindered normalized total fraction, RHNT: right hindered normalized total fraction, LFNI: left free normalized isotropic fraction, RFNI: right free normalized isotropic fraction.

## Methods

### Participants and ABCD Study Design

We used cross-sectional data from the United States’ ABCD Study^®^ annual 5.1 data release (2023) (doi:10.15154/8873-zj65, NDA Study; n = 11,868), including sociodemographic, neuroimaging, and linked external environmental data. The ABCD Study originally enrolled 11,880 children (between 2016 - 2018) aged 9 to 10 years from 21 sites across the United States and is following them longitudinally for ten years ^51^. The ABCD Study sample mirrors typical U.S. distributions for age, sex, and household size, but it underrepresents children of Asian, American Indian/Alaskan Native, and Native Hawaiian/Pacific Islander ancestry ^62^. Participants were excluded if they lacked English proficiency, had severe sensory, neurological, medical, or intellectual limitations, or were unable to complete an MRI scan. The ABCD study received approval from the Institutional Review Board (IRB) and Human Research Protections Programs (HRRP) at the University of California, San Diego. The IRB at the University of California, San Diego oversees all experimental and consent procedures with IRB board approval at the 21 ABCD sites. Written assent was provided by each participant, with written consent obtained from their legal guardians. For further details, see ^51,52^. All analyses described below, apart from the hippocampal long-axis volume data (see below), used data pre-processed and tabulated by the Data Analysis, Informatics, and Resource Center (DAIRC) at the University of California, San Diego. In this study, we analyzed a subset of the ABCD Study dataset (Supplemental Figure 1), applying exclusion criteria based on MRI quality, incidental findings, and the availability of complete case data. This process resulted in a sample of 7,940 participants for our RSI analyses. To ensure consistency, we matched participants with available long-axis subregion data to those in the RSI analyses, yielding a sample of 6,795 participants.

### Residential Ambient Air Pollution Estimates and PM_2.5_ Sources

We assigned 2016 annual average air pollution exposure to each child’s primary residence collected from the caregiver during the initial ABCD Study enrollment visits (2016-2018)^63^. Using hybrid spatiotemporal models combining land-use regression, satellite-based aerosol optical depth, and chemical transport were used to derive daily estimates of PM_2.5_ (µg/m^3^), NO_2_ (parts per billion [ppb]), and 8-hour maximum ground-level O_3_ (ppb) at a 1-km^2^ resolution ^64–66^. These hybrid spatiotemporal models show excellent model performance, showing high out-of-sample validation (i.e., model R^2^ based on 10-fold cross-validations: PM_2.5_ R^2^=0.86; NO_2_ R^2^=0.79; O_3_ R^2^=0.90) ^64–66^ To estimate residential exposure to PM_2.5_ constituent components, similar machine learning-based models were used to calculate 15 components, including ammonium (NH_4_^+^), bromine (Br), calcium (Ca), copper (Cu), elemental carbon (EC), iron (Fe), lead (Pb), nickel (Ni), nitrate (NO_3−_), organic carbon (OC), potassium (K), silicon (Si), sulfate (SO_4_^2-^), vanadium (V), and zinc (Zn) ^54,67–69^. We selected these components based on sufficient monitoring data and that these components may exhibit varying degrees of toxicity that are important to capture ^54,70,71^. Although trace elements comprise a small fraction of PM_2.5_ total mass, evidence indicates they play a significant role in PM_2.5_ neurotoxicity ^2,37,72^. Trace elements (particularly redox-active metals such as Fe and Cu) can increase oxidative potential and promote oxidative stress, neuroinflammation, and metal dyshomeostasis-mechanisms implicated in PM_2.5_-related neurotoxicity ^15,37^. Consistent with this, PM toxicity varies by chemical composition and emission source, motivating component- and source-resolved exposure analyses ^14^.

Monthly estimates for each component were calculated at a spatial resolution of 50-m^2^ accounting for 166 predictors, including seasonality, geographical distribution, meteorological data, and proximity to power plants and highways ^54,67–69^. The model performance for the components were also high, with out-of-sample validation R^2^ for individual components ranging from 0.80 (Cu) to 0.96 (SO_4_^2-^) across the fifteen different constituents.

Based on our previously published work ^10^, we also estimated for each individual exposure estimates to six source factors, including crustal, ammonium sulfates, biomass burning, ammonium nitrates, traffic, as well as industrial and residual oil burning (Figure 1A). The exact methods of how these six sources were derived for the ABCD Study cohort have been previously published in detail ^10^. Briefly, to obtain PM_2.5_ source factors of major emission sources and quantify their contribution to PM_2.5_, positive matrix factorization (PMF) was applied to individual participant-level PM_2.5_ component estimates for ABCD participants using the PMF tool developed by the EPA (EPA, v5.0)^73^. Model inputs included component concentrations and uncertainty estimates based on the root mean square error from spatiotemporal prediction models, and multiple solutions (4-8 factors) were evaluated using rotational and displacement analyses to select the most statistically robust source profiles ^10^. Through these analyses, PMF analysis decomposed the fifteen components into six factors, which were conceptualized into potential sources based on expert knowledge regarding the components that largely loaded onto those factors and the geographic trends in their contributions ^10^. The six sources included: crustal materials (driven largely by Si, Ca), ammonium sulfates (driven largely by NH_4_^+^, SO_4_^2-^), biomass burning (driven largely by OC, K, Br), traffic emissions (driven largely by EC, OC, Fe, Cu), ammonium nitrates (driven largely by NH_4_^+^, NO_3_^-^), and industrial/residual fuel burning (driven largely by Pb, Zn, Ni) (Figure 1A) ^10^. For interpretative and applied purposes, these sources are defined largely by their dominant components; however, it is important to note other components contribute to each source’s overall composition (Figure 1A).

As expected, PM_2.5_ components are generally positively correlated with each other (*ρ* = −0.07 to 0.83). O_3_ showed negative correlations with most pollutants (*ρ* = −0.32 to 0), except for traffic and copper, with which it was positively correlated. NO_2_ was mostly positively correlated with other pollutants (*ρ* = 0.01 to 0.56), except for SO_4_^2-^ and secondary ammonium sulfates. The various sources were strongly to moderately interrelated, with correlations ranging from *ρ* = −0.49 to 0.25 (Supplemental Figures 2-3).

### Brain MRI Acquisition and Processing

Structural MRI protocols for the ABCD study have been extensively described ^74,75^. Briefly, a harmonized protocol was used across sites with either a Siemens, Phillips, or GE 3T MRI scanner. Motion compliance training and real-time, prospective motion correction were employed to reduce motion distortion. T1-weighted (TE 2–2.9 ms, TR 6.31–2500 ms, T1 1060 ms, flip angle 8 degrees, FOV 256 x 256, resolution 1mm³ isotropic, 176 slices) and T2-weighted images (TE 60–565 ms, FOV 256 x 256, resolution 1mm³ isotropic, 176 slices) were acquired using a magnetization-prepared rapid acquisition gradient echo (MPRAGE) and fast spin echo sequence with variable flip angle, respectively ^74^. The DWI acquisition protocols for the ABCD Study included a voxel size of 1.7 mm isotropic and employed multiband echo planar imaging ^76,77^ with a slice acceleration factor of 3. Each DWI acquisition included a fieldmap scan for B0 distortion correction. The ABCD Study used a multi-shell diffusion acquisition protocol that included 7 b=0 frames and 96 total diffusion directions at 4 b-values (6 with b = 500 s/mm², 15 with b = 1000 s/mm², 15 with b = 2000 s/mm², and 60 with b = 3000 s/mm²). All images underwent distortion correction, bias field correction, motion correction, and manual and automated quality control ^75^. Only images without clinically significant incidental findings (*mrif_score = 1*) that passed all ABCD Study quality-control parameters (*imgincl_dmri_include = 1*) were included in analyses. Moreover, based on previous work showing PM_2.5_ associated hemispheric differences in brain structure ^31,78,79^, we chose to examine hippocampal structure in each hemisphere.

RSI, a multicompartmental biophysical diffusion model was implemented using all 96 DWI directions collected. RSI extends beyond conventional approaches such as Diffusion Tensor Imaging and Neurite Orientation Dispersion and Density Imaging (NODDI) by modeling multiple tissue compartments—intracellular, extracellular, and free water—to capture a broader spectrum of water diffusion behavior with greater microstructural specificity ^55–58^. Unlike NODDI, which may overestimate neurite density in regions with complex tissue architecture ^80^ and provides information only about intracellular diffusion ^35,81^, RSI provides detailed information about the volume-fractions of both intracellular and extracellular tissue compartments of the brain ^57,58^. To date, RSI has been used in neuroradiological applications to estimate cellular density in brain tumors by distinguishing diffusion signals arising from both intracellular and extracellular environments ^57,58^. Moreover, RSI has been histologically validated in neuro-oncological applications ^58^ and in subcortical structures that contain mixed cellular and myelinated compartments, similar to the hippocampus, such as the striatum and globus pallidus, highlighting its sensitivity to biologically meaningful microstructural variation ^57^. In addition to neuro-radiological clinical applications ^57,58^, RSI has been increasingly used in developmental and environmental neuroimaging studies to characterize both cross-sectional and longitudinal differences in gray and white matter microstructure in relation to air pollution exposure ^48,53,82–84^, environmental stress exposure ^85^, and lower socioeconomic status ^86^. RSI model outputs include normalized measures, all of which are unitless on a scale of 0 to 1 and represent signal fractions of the diffusion signal. Restricted normalized total signal fraction (RNT) indicates total intracellular diffusion within a voxel, where an increase may represent myelination, dendritic sprouting, arborization, or increases in neurite density ^55^. Hindered normalized total signal fraction (HNT) measures total extracellular diffusion and is inversely related to RNT; increased HNT values may represent demyelination, an increased number of mature astrocytes, an increase in cell body size, or fewer cells and neurites ^55^. Free normalized isotropic signal fraction (FNI) represents free water movement and cerebrospinal fluid ^55,57^. In RSI, of the signal fractions, the restricted component may be interpreted as primarily intra-neurite water (inside cells/axons/dendrites), whereas the hindered component may reflect primarily extra-neurite water (extracellular spaces) ^57,58^. Thus, greater hindered and lower restricted signals may reflect proportionally higher extracellular signal and lower intracellular signal of the diffusion signal. In general, higher RNT values reflect proportionally greater intracellular signal fractions and may indicate greater cellular density or neurite complexity, whereas higher HNT and FNI values reflect proportionally greater extracellular and free water signal contributions, respectively ^56–58^. However, RSI metrics represent relative signal fractions rather than direct histological measures and should be interpreted as indicators of tissue microstructural composition rather than definitive markers of tissue integrity ^57,58^. In the current study, RSI analyses focused on the hippocampal region of interest, which was derived from FreeSurfer’s automated, atlas-based, volumetric segmentation procedure (*aseg*). RSI measures were then collated by hemisphere resulting in six RSI metrics, with left and right RNT, HNT, and FNI diffusion signal fractions used in our analyses (Figure 1B).

### Hippocampal Long-Axis Subregion Processing

Segmentation of hippocampal longitudinal subregions was conducted using both T1-weighted and T2-weighted images by co-author J.G.P. with the Freesurfer v7.0 automated hippocampal subregion segmentation tool ^59,60^. This tool uses a probabilistic atlas derived from ultra-high-resolution (7T) ex vivo MRI data (0.13 mm^3^) to identify hippocampal subregions (head, body, and tail) and subfields (e.g., dentate gyrus, subiculum), followed by analysis using in vivo T1-weighted (1mm^3^) to distinguish the hippocampus from surrounding brain regions ^60^. Methods for these analyses are described in more detail at: https://collection3165.readthedocs.io/en/stable/. A subset of these hippocampal subregion data has been previously used to determine hippocampal genetic correlates along its long-axis ^61^. Here, we examine left and right hippocampal long-axis volumes (head, body, and tail) (mm^3^) (Figure 1C).

### Rey Auditory Verbal Learning Test (RAVLT)

Learning and memory were measured using the Rey Auditory Verbal Learning Test (RAVLT)^87,88^. The RAVLT evaluates verbal memory by presenting a list of 15 unrelated words (List A) over five learning trials, with participants recalling as many words as possible after each trial. Following these trials, a second list of 15 words (List B) is introduced as a proactive mnemonic interference task, after which participants immediately recall List A. Delayed recall of List A was then assessed 20–30 minutes later. Our primary outcomes included: learning (the difference in words recalled between Trial 5 and Trial 1), proactive mnemonic interference (total words recalled from List B), immediate recall (total words recalled from List A immediately after interference), and delayed recall (total words recalled from List A after the delay). On average, participants learned approximately 6 words across Trials 1 through 5, remembered approximately 9.8 words during immediate recall, 9.3 words during delayed recall, and 4.9 during proactive interference (List B) (Supplemental Table 6).

### Statistical Analyses

We conducted all statistical analyses using R(v.4.4.0) and used the ExPosition and TExPosition packages ^89^. We examined the relationships between air pollution exposure and hippocampal metrics using a series of Partial Least Squares Correlations (PLSC)^90,91^. PLSC is a multivariate technique that identifies latent patterns of covariance between interrelated sets of variables. Thus, this method is particularly well-suited for our study, given the high intercorrelations among our exposure, brain, and cognitive variables of interest (Supplementary Figures 2-5).

PLSC analyses require independent and complete-case data. Thus, before analysis, we cleaned the datasets to remove missing values and specific participants flagged during initial quality control (*imgincl_dmri_include = 1*, *imgincl_t1w_include = 1, mrif_score = 1 or 2* which reflect no abnormal MRI findings; *not missing family ID*) resulting in 9,199 participants for selected variables of interest, including covariates (see below) and one child per family. Missing data across selected variables ranged from 0.008% to 5.74% (Supplemental Table 2), with the largest proportion attributable to missing address information, which precluded estimation of air pollution exposure levels. Previous research from our lab has demonstrated that missing address data within the ABCD Study are likely missing at random ^46^. Given the relatively low overall percentage of missingness and the absence of evidence suggesting systematic bias, we employed listwise deletion to retain a consistent analytic sample across models. This resulted in a final sample of 7,940 participants for the microstructure analyses. Next, we selected a subset of these participants who had data available for hippocampal long-axis volume, for a final sample of 6,795 for these analyses (Supplemental Figure 1; Table 1; Supplemental Table 1; Supplemental Table 3).

**Table 1.** Sample participant characteristics.

|  | Hippocampal RSI<br>(N=7940) | Hippocampal Long-Axis Volume<br>(N=6795) |
| --- | --- | --- |
| **Age (months)** |  |  |
| Mean (SD) | 120 (± 7.4) | 120 (± 7.4) |
| **Assigned Sex at Birth** |  |  |
| Female | 4114 (52 %) | 3512 (52 %) |
| Male | 3826 (48 %) | 3283 (48 %) |
| **Race & Ethnicity** |  |  |
| non-Hispanic white | 4162 (52 %) | 3719 (55 %) |
| non-Hispanic Black | 1110 (14 %) | 882 (13 %) |
| Hispanic | 1676 (21 %) | 1362 (20 %) |
| Asian/Other<sup>a</sup> | 992 (12 %) | 832 (12 %) |
| **Overall Income** |  |  |
| <\$50K | 2108 (27 %) | 1739 (26 %) |
| ≥\$50K and <\$100K | 2065 (26 %) | 1826 (27 %) |
| ≥\$100K | 3103 (39 %) | 2680 (39 %) |
| Don't know/Refuse to answer | 664 (8 %) | 550 (8 %) |
| **Parental Highest Education** |  |  |
| < HS Diploma | 382 (5 %) | 272 (4 %) |
| HS Diploma/GED | 729 (9 %) | 574 (8 %) |
| Some College | 2041 (26 %) | 1762 (26 %) |
| Bachelor's Degree | 1999 (25 %) | 1755 (26 %) |
| Post Graduate Degree | 2781 (35 %) | 2425 (36 %) |
| Missing/Refused | <10 (<1 %) | <10 (<1 %) |
| **MRI Manufacturer** |  |  |
| GE Medical Systems | 1870 (24 %) | 1465 (22 %) |
| Philips Medical Systems | 891 (11 %) | 679 (10 %) |
| SIEMENS | 5179 (65 %) | 4651 (68 %) |
| **Urban vs Rural** |  |  |
| Urbanized Area | 7036 (89 %) | 5978 (88 %) |
| Urban Clusters | 242 (3 %) | 221 (3 %) |
| Rural | 662 (8 %) | 596 (9 %) |
| **Neighborhood Safety** |  |  |
| Mean (SD) | 3.9 (± 0.97) | 3.9 (± 0.95) |

<sup>a</sup> Race/ethnicity category includes participants identified by their caregiver as American Indian/Native American, Alaska Native, Native Hawaiian, Guamanian, Samoan, Other Pacific Islander, Asian Indian, Chinese, Filipino, Japanese, Korean, Vietnamese, Other Asian not listed, or Other Race not listed.

Using these final sample sizes, we then applied multiple linear regression models to obtain residualized hippocampal, air pollution, and learning and memory variables, accounting for potential confounding, using the *residuals()* function in *R*. By using the residuals from our linear models we controlled for the influence of demographic, socioeconomic, and neuroimaging covariates, thereby isolating the specific relationships between the brain imaging and cognitive variables and air pollution exposure. Specifically, to account for the higher pollution levels often found among structurally disadvantaged groups ^92,93^, we accounted for race/ethnicity (*non-Hispanic White, non-Hispanic Black, Hispanic/Latino,* or *Asian/Other*), total household income in USD (≥*100K,* ≥*50-<100K, <50K,* or *Don’t Know/Refuse to Answe*r), highest household education (*Post-Graduate, Bachelor, Some College, High School Diploma/GED,* or *<High School Diploma*), proximity to major roadways in meters, urbanicity (urban area, urban cluster, rural)^94^. and PhenX parent-report of neighborhood safety^95^ (Table 1). We also controlled for individual differences by including each child’s parent-reported sex assigned at birth (*male* or *female*), handedness *(right, left,* or *mixed*), and age (in months). For our microstructure and long-axis volume analyses, we further controlled for diffusion imaging mean motion (as indexed by framewise displacement) and intracranial volume, respectively. To mitigate variability introduced by imaging equipment we included MRI scanner serial number (n = 29) The residuals then served as the input for our subsequent PLSC analyses.

### Partial Least Squares Correlation (PLSC)

We applied a series of PLSC analyses to examine the multivariate relationship between residualized 1) air pollution data, 2) hippocampal microstructure, 3) long-axis volumes, and 4) RAVLT performance accounting for confounds and precision variables. PLSC allows us to identify latent dimensions (hidden patterns) that maximize the covariance between two sets of matrices, or variables (Figure 1D) ^90,91,96^. PLSC uses Singular Value Decomposition (SVD) to break down the correlation between the two sets of variables into singular values and saliences (akin to variable loadings in Principal Component Analysis). While singular values capture the strength of the amount of shared covariance, saliences represent the contribution of original variables towards the latent dimensions ^90,96^. In contrast to the traditional use of independent and dependent variables in analyses such as regression, variables included in PLSC are blocked as a set of variables ^90,91,96^. For example, as in our first PLSC analysis, we included PM_2.5_, NO_2_, and O_3_ in one set (***X***-set) and hippocampal microstructure (i.e., RNT, HNT, and FNI) in another set (***Y***-set) (Figure 1D). The singular values and loadings are characteristics of each latent dimension that capture associations between the two sets by analyzing the information common between them ^90,91,96^. In other words, PLSC latent dimensions capture the shared covariance structure between the two sets of variables in the matrices. This differs from univariate regression, where variance explained refers to the percent of the outcome explained by an independent variable. Positive and negative loadings reflect opposing patterns of covariance within a latent dimension and should not be interpreted as directional or causal effects. Instead, loadings indicate the relative contribution of each variable to the multivariate covariance structure captured by the latent dimension and must be interpreted as part of an integrated multivariate pattern rather than as independent associations between individual variables.

To determine if the overall PLSC model is significant, permutation tests with 10,000 iterations were conducted and an omnibus p value was calculated as the proportion of iterations where the sum of eigenvalues from all latent dimensions exceeded the observed sum of eigenvalues (i.e., that is, the total variance explained by the observed latent dimensions exceeded that expected under permutation). In addition to the overall model significance, the significance of each individual latent dimension was also evaluated as the proportion of iterations where the permuted eigenvalue of that latent dimension exceeded its observed eigenvalue. In PLSC, the maximum number of extractable latent dimensions is capped by the number of variables in the smaller block (e.g., three dimensions for models pairing three air pollution exposures with six hippocampal variables). Of the max number of dimensions, only latent dimensions with a permutation-derived p-value < .05 over 10,000 tests were retained for interpretation. This threshold was applied consistently across all PLSC analyses, meaning that in some models only the first latent dimension met this criterion, while in others more than one did. Figures display only retained (statistically significant) latent dimensions; variation in the number of dimensions shown across figures therefore directly reflects variation in the number of significant dimensions identified in each model. The significance of each latent dimension is then determined by comparing the observed singular value to the distribution of permuted singular values, with the p-value reflecting the proportion of permutations in which the permuted singular value exceeded the observed value. Latent dimensions with p < .05 under this permutation test were considered statistically significant and retained; non-significant dimensions were extracted but not interpreted or displayed. Note that in some models, only the first latent dimension met the significance criterion, while others had multiple significant dimensions.

To assess the stability of the identified latent dimensions, we employed bootstrapping techniques resampling 10,000 times, leaving one sample out each time (with the help of *Boot4PLSC()* function in *R*)^97^ and recalculate the PLSC components to generate bootstrapped ratios for each variable. This procedure enabled us to rigorously evaluate both the overall importance of each latent dimension and the stability of individual variable contributions ^91^. Variables with ratios exceeding a threshold of ±2.5 were considered significant (equivalent to *p* < .01) contributors to the latent dimensions. In PLSC, permutation testing and bootstrapping provide complementary safeguards against false positives. Permutation testing evaluates whether the latent dimensions reliably distinguish signal from noise, whereas bootstrap ratios (calculated as the weight determined by SVD analysis divided by the bootstrap-estimated standard error) scores stability of individual variable contributions ^91,98,99^.

Using this PLSC approach, we conducted five separate sets of PLSCs examining: (1) PM_2.5_ total mass, NO_2_, and O_3_ with hippocampal structure (i.e., RSI outcomes or long-axis volumes); (2) fifteen PM_2.5_ chemical components (alongside NO_2_ and O_3_) with hippocampal structure (i.e., RSI outcomes or long-axis volumes); (3) six PM_2.5_ source factors (i.e., crustal, ammonium nitrates, ammonium sulfates, biomass burning, traffic, industrial/residual fuel burning) with hippocampal structure (i.e., RSI outcomes or longitudinal-axis volumes); (4) air pollutants with RAVLT performance; and (5) hippocampal structure (i.e., RSI outcomes or longitudinal-axis volumes) with RAVLT performance. For each analysis, permutation test results, including the total number of dimensions tested and the number retained, are reported in the corresponding Supplemental Tables 7-10. Figures 2-5 display only the retained, statistically significant latent dimensions for each model, and dimensions are labeled consistently (e.g., Dimension 1, Dimension 2) across all figures.

**Figure 2.**
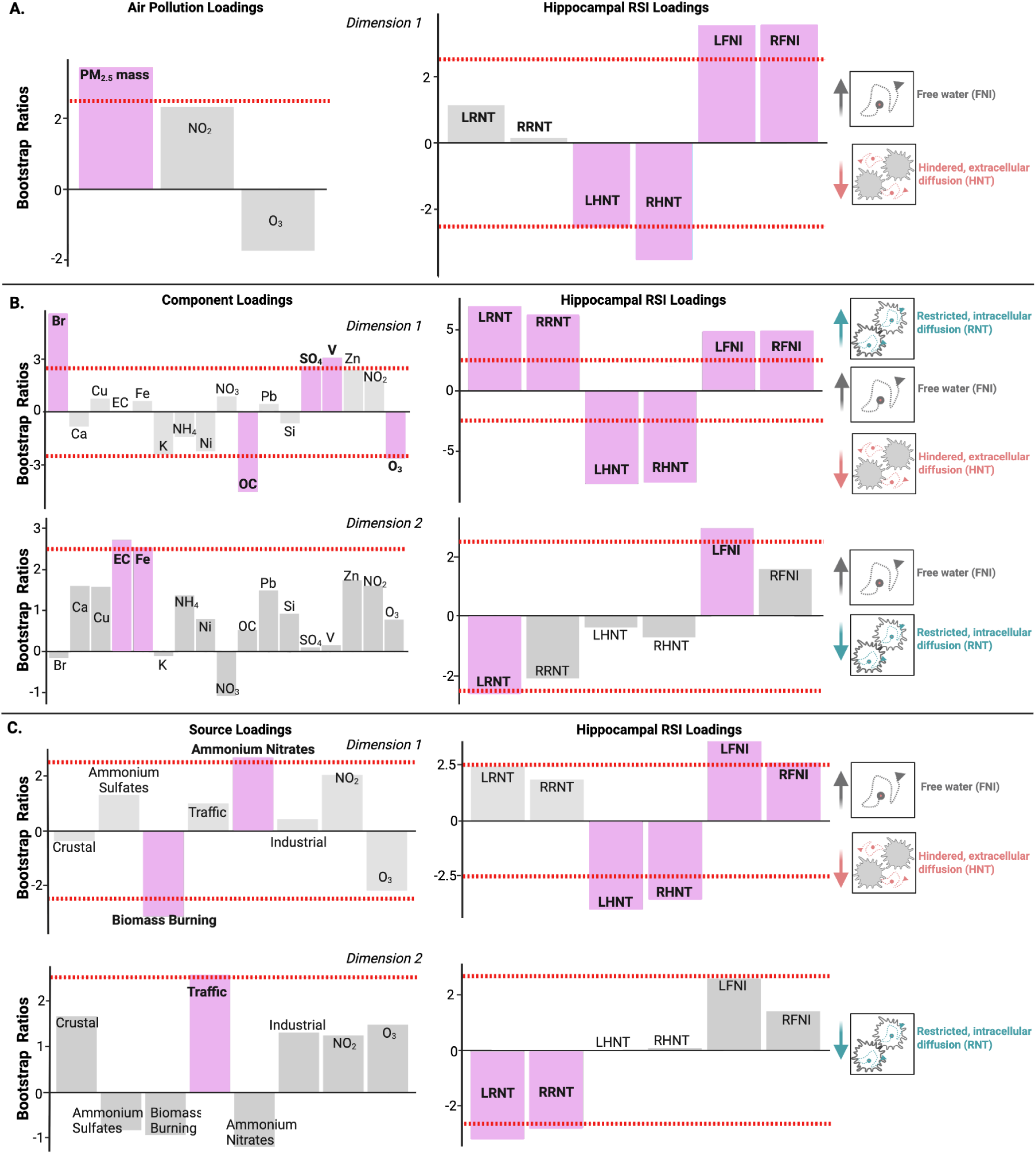
Association between air pollution and hippocampal microstructure at 9-11 years (n = 7,940). Permutation testing identified statistically significant latent dimensions linking air pollution exposures and hippocampal RSI metrics; only significant latent dimensions are shown (Supplemental Figures 6-8; Supplemental Table 7). Bars represent bootstrap ratios, with dashed lines at +2.5 indicating stability thresholds (approximately *p* < .01). Positive and negative bootstrap ratios reflect opposing patterns of covariance within each latent dimension. **A.** Higher annual average PM_2.5_ total mass exposure is associated with higher bilateral FNI and lower bilateral HNT diffusion. **B.** In the first latent dimension, PM_2.5_ components based analyses showed higher Br, SO_4_, and V, but lower OC and O_3_ exposure is associated with lower bilateral HNT as well as higher bilateral FNI and RNT diffusion. In the second latent dimension, higher EC and Fe exposure is associated with lower LRNT and higher LFNI. **C.** PM_2.5_ source analyses showed that for the first latent dimension, higher ammonium nitrate but lower biomass burning exposure is associated with lower bilateral HNT and higher bilateral FNI. whereas for the second latent dimension higher traffic exposure is associated with lower bilateral RNT. respectively. Abbreviations: Br: bromine. Ca: calcium, Cu: copper, EC: elemental carbon. Fe: iron, K: potassium, NH_4_: ammonium. Ni: nickel, NO_2_: nitrogen dioxide, NO_3_: nitrate, O_3_: ozone, OC: organic carbon, Pb: lead, PM_2.5_ mass: fine particulate matter. Si: silicon, SO_4_: sulfate, V: vanadium, Zn: zinc, RSI: restriction spectrum imaging; LRNT: left restricted (intracellular) normalized total fraction, RRNT: right restricted (intracellular) normalized total fraction, LHNT: left hindered (extracellular) normalized total fraction, RHNT: right hindered (extracellular) normalized total fraction, LFNI: left free (water) normalized isotropic fraction, RFNI: right free (water) normalized isotropic fraction.

## Results

The characteristics of the participant sample (n = 7,940 for microstructure and n = 6,795 for long axis volume) are detailed in Table 1. A comparison between our analytic sample and the broader ABCD Study cohort is provided in Supplemental Table 1. Participants are similar in terms of age, sex, race/ethnicity, and parental education and income. The descriptive statistics of the hippocampal RSI microstructure values and hippocampal long-axis volume estimates are reported in Supplemental Table 4 whereas the air pollution exposure estimates for the study sample are reported in Supplemental Table 5. To contextualize exposure levels, one-sample *t*-tests were conducted to compare mean concentrations of PM_2.5_, NO_2_, and O_3_ in the current sample to current 2024 EPA standards ^11,100^ and 2021 World Health Organization recommendations ^101^. Mean pollutant concentrations across participants (n = 7,940) were 7.7 ± 1.6 µg/m^3^ for PM_2.5_, 18.7 ± 5.7 ppb for NO_2_, 41.7 ± 4.4 ppb for O_3_. Relative to the EPA standards (PM_2.5_ = 9 µg/m^3^; NO_2_ = 53 ppb; O_3_ 8-hour maximum = 70 ppb), PM_2.5_, NO_2_, and O_3_ levels were significantly lower (PM_2.5_: *t*(7939) = −75.11, *p* < 0.001; NO_2_: *t*(7939) = −539.65, *p* < 0.001; O_3_: (*t*(7939) = −569.10, *p* < 0.001). When compared with the more stringent WHO 2021 Air Quality Guidelines (PM_2.5_ = 5 µg/m^3^; NO_2_ = 10 ppb; O_3_ = 60 ppb, study sample concentrations were significantly higher for PM_2.5_ (*t*(7939) = 150.66, *p* < 0.001) and NO_2_ (*t*(7939) = 136.26, *p* < 0.001), but significantly lower for O_3_: *t*(7939) = −368.18, *p* < 0.001).

### Air pollution and hippocampal microstructure

For the model including annual average PM_2.5_ total mass, NO_2_, and O_3_, permutation testing indicated that the overall PLSC model was statistically significant (*p* = .0001; Supplemental Table 7). Of the three latent dimensions, only the first latent dimension exceeded the permutation-derived significance threshold, while subsequent latent dimensions were not significant (Supplemental Figure 6). Therefore, only the first latent dimension was retained for interpretation. This latent dimension accounted for 81% of the covariance captured by the model and showed that higher annual average PM_2.5_ total mass was associated with higher bilateral free water (LFNI and RFNI) and lower bilateral hindered, extracellular diffusion (LHNT and RHNT) (Figure 2A). Next, we examined associations between fifteen PM_2.5_ chemical components (alongside NO_2_ and O_3_) and hippocampal RSI metrics. Permutation testing indicated that the overall PLSC model was statistically significant (*p* = .0001; Supplemental Table 7). Of the six latent dimensions, the first and second latent dimensions exceeded the permutation-derived significance threshold, while subsequent latent dimensions were not significant (Supplemental Figure 7). These two latent dimensions accounted for 72% and 24% of the covariance captured by the model, respectively. In the first latent dimension, higher annual concentrations of Br, SO_4_^2-^, and V, but lower levels of OC and O_3_, were associated with lower bilateral hindered, extracellular diffusion (HNT) as well as higher bilateral restricted, intracellular (RNT) and free water (FNI) diffusion (Figure 2B). In the second latent dimension, higher annual exposure to EC and Fe was associated with lower left hemisphere restricted, intracellular diffusion (RNT) and higher left hemisphere free water (FNI) diffusion (Figure 2B). We observed a similar pattern for PM_2.5_ source factors and hippocampal microstructure. Permutation testing indicated that the overall PLSC model was statistically significant (*p* = .0001; Supplemental Table 7). Of the six latent dimensions, the first and second latent dimensions exceeded the permutation-derived significance threshold, while subsequent latent dimensions were not significant (Supplemental Figure 8). These two latent dimensions accounted for 61% and 32% of the covariance captured by the model, respectively. In the first latent dimension, higher annual exposure to ammonium nitrates, but lower exposure to biomass burning, was associated with lower bilateral hindered, extracellular diffusion (HNT) and higher bilateral free water (FNI). In the second latent dimension, higher annual levels of traffic-related exposure were associated with lower bilateral restricted, intracellular diffusion (RNT) (Figure 2C).

### Air pollution and volumes of the hippocampal head, body, and tail

For the model including annual average PM_2.5_ total mass, NO_2_, and O_3_ and hippocampal head, body, and tail, permutation testing indicated that the overall PLSC model was not statistically significant (*p* = .98; Supplemental Table 8; Supplemental Figure 9). In contrast, permutation testing indicated that the overall PLSC model examining fifteen PM_2.5_ components (alongside NO_2_ and O_3_) and hippocampal long-axis volumes was statistically significant (*p* = .0004; Supplemental Table 8). Of the six latent dimensions, only the first latent dimension exceeded the permutation-derived significance threshold, while subsequent latent dimensions were not significant (Supplemental Figure 10). The first latent dimension accounted for 75% of the covariance captured by the model and showed that higher annual exposure to Cu and Zn was associated with smaller hippocampal volumes of the left head and the right body and tail (Figure 3A). Similarly, permutation testing indicated that the overall PLSC model examining PM_2.5_ source factors and hippocampal long-axis volumes was statistically significant (*p* = .041; Supplemental Table 8). Of the six latent dimensions, only the first latent dimension exceeded the permutation-derived significance threshold, while subsequent latent dimensions were not significant (Supplemental Figure 11). The retained latent dimension accounted for 77% of the covariance captured by the model and showed that higher annual levels of traffic- and industrial-related air pollution exposure were associated with smaller hippocampal volumes of the left head and body and the right body and tail (Figure 3B).

**Figure 3.**
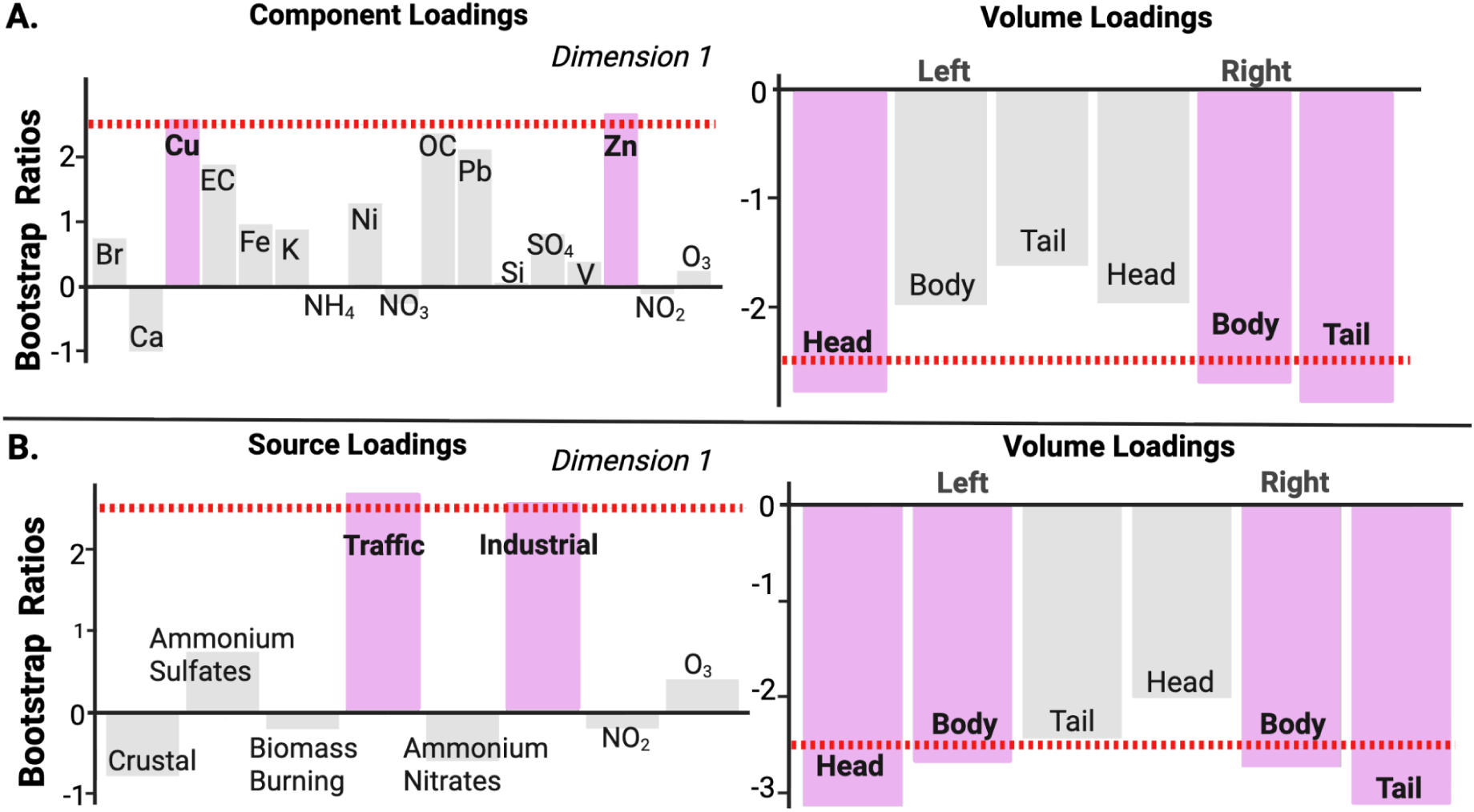
Association between air pollution and hippocampal long-axis volume at 9-11 years (n = 6,795). Permutation testing identified statistically significant latent dimensions linking air pollution exposures and hippocampal long-axis volumes: only significant latent dimensions are shown (Supplemental Figures 9-11; Supplemental Table 8). Bars represent bootstrap ratios, with dashed lines at ±2.5 indicating stability thresholds (approximately p < .01). Positive and negative bootstrap ratios reflect opposing patterns of covariance within each latent dimension. **A.** PM_2.5_ components based analyses showed higher Cu and Zn exposure is associated with smaller left hippocampal head as well as the right hippocampal body and tail volume. B. PM_2.5_ source analyses showed traffic and industrial-related emission exposure is associated with smaller left hippocampal head and body as well as the right hippocampal body and tail volumes. Abbreviations: Br: bromine, Ca: calcium, Cu: copper, EC: elemental carbon, Fe: iron, K: potassium, NH_4_: ammonium, Ni: nickel, NO_2_: nitrogen dioxide, NO_3_: nitrate, O_3_: ozone, OC: organic carbon, Pb: lead, PM_2.5_: fine particulate matter, Si: silicon, SO_4_: sulfate, V: vanadium, Zn: zinc.

### Air pollution and RAVLT performance

For the model including annual average PM_2.5_ total mass, NO_2_, and O_3_, permutation testing indicated that the overall PLSC model was not statistically significant (*p* = .96; Supplemental Table 9;Supplemental Figure 12). In contrast, permutation testing indicated that the overall PLSC model examining fifteen PM_2.5_ chemical components (alongside NO_2_ and O_3_) and RAVLT performance was statistically significant (*p* = .0001; Supplemental Table 9). Of the four latent dimensions,only the first latent dimension exceeded the permutation-derived significance threshold, while subsequent latent dimensions were not significant (Supplemental Figure 13). The retained latent dimension accounted for 66.5% of the covariance captured by the model and showed that higher annual concentrations of Ca, EC, and Zn were associated with poorer learning and immediate recall performance (Figure 4). Similarly, permutation testing indicated that the overall PLSC model examining PM_2.5_ source factors and RAVLT performance was statistically significant (*p* = .002; Supplemental Table 9). Of the four latent dimensions, the first and second latent dimensions exceeded the permutation-derived significance threshold, while subsequent latent dimensions were not significant (Supplemental Figure 14). These two latent dimensions accounted for 58.24% and 38.18% of the covariance captured by the model, respectively, and reflected associations between air pollution source exposure patterns and poorer learning and immediate recall performance. However, bootstrap resampling indicated limited stability of individual variable contributions within these latent dimensions (Supplemental Figure 15), suggesting caution in interpreting the relative importance of specific sources while supporting the robustness of the overall multivariate covariance pattern.

**Figure 4.**
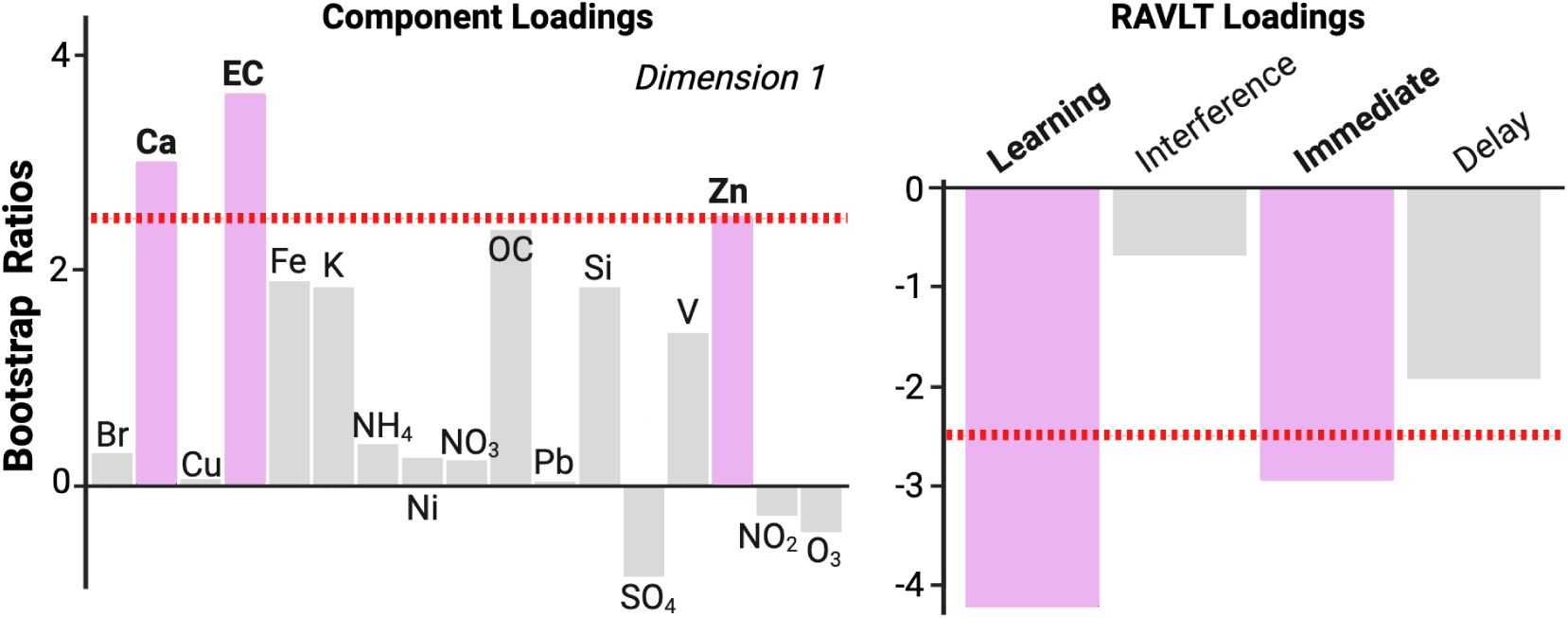
Associations between air pollution exposure and RAVLT performance at 9-11 years (n = 7,699). Permutation testing identified statistically significant latent dimensions linking air pollution exposures and RAVLT performance; only significant latent dimensions are shown (Supplemental Figures 12-14; Supplemental Table 9). Bars represent bootstrap ratios, with dashed lines at ±2.5 indicating stability thresholds (approximately p < .01). Positive and negative bootstrap ratios reflect opposing patterns of covariance within each latent dimension. Higher Ca, EC, and Zn exposure is associated with poorer learning and immediate recall performance. For the RAVLT, learning = the difference in words recalled between Trial 5 and Trial 1 from List A; interference = proactive mnemonic interference as measured by total words recalled from List B; immediate recall = total words recalled from List A immediately after interference; delayed recall= total words recalled from List A after the delay. Abbreviations: Br: bromine, Ca: calcium, Cu: copper, EC: elemental carbon, Fe: iron, K: potassium, NH_4_: ammonium, Ni: nickel, NO_2_: nitrogen dioxide, NO_3_: nitrate, O_3_: ozone, OC: organic carbon, Pb: lead, Si: silicon, SO_4_: sulfate, V: vanadium, Zn: zinc, RAVLT: Rey Auditory Verbal

### Hippocampal Structure and RAVLT performance

The PLSC analysis examining associations between hippocampal diffusion metrics and RAVLT performance revealed that permutation testing indicated the overall model was statistically significant (*p* = .0001; Supplemental Table 10). Of the four latent dimensions, only the first latent dimension exceeded the permutation-derived significance threshold, while subsequent latent dimensions were not significant (Supplemental Figure 16). The retained latent dimension accounted for 82% of the covariance captured by the model and showed that higher left hemisphere free water (FNI), alongside lower left hemisphere hindered, extracellular diffusion (HNT), was associated with poorer immediate and delayed recall performance. Similarly, permutation testing indicated that the overall PLSC model examining hippocampal long-axis volumes and RAVLT performance was statistically significant (*p* = .0001; Supplemental Table 10). Of the four latent dimensions, only the first latent dimension exceeded the permutation-derived significance threshold, while subsequent latent dimensions were not significant (Supplemental Figure 17). The retained latent dimension accounted for 96% of the covariance captured by the model and showed that smaller bilateral hippocampal tail volumes were associated with poorer immediate recall performance (Figure 5B).

**Figure 5.**
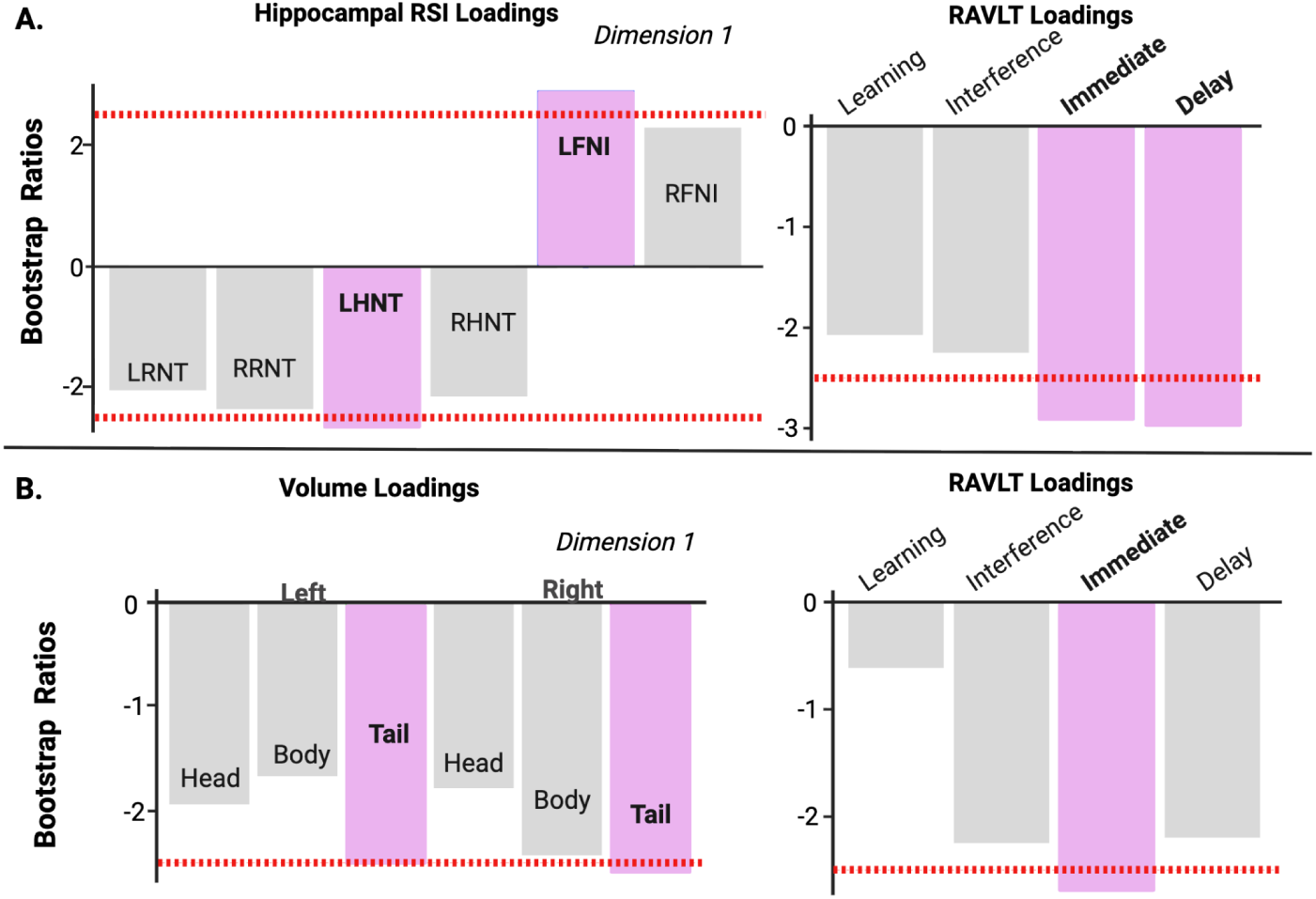
Associations between hippocampal architecture and RAVLT performance at 9-11 years. Permutation testing identified statistically significant latent dimensions linking hippocampal diffusion metrics and long-axis volumes with RAVLT performance: only significant latent dimensions are shown (Supplemental Figures 16—17; Supplemental Table 10). Bars represent bootstrap ratios, with dashed lines at +2.5 indicating stability thresholds (approximately *p <* .01). Positive and negative bootstrap ratios reflect opposing patterns of covariance within each latent dimension. **A.** Higher left free water alongside lower left hindered waler diffusion is linked to poorer immediate and delayed recall performance (n = 7,699). **B.** Lower bilateral hippocampal tail volumes are linked to poorer immediate memory performance (n = 6,589). For the RAVLT, learning = the difference in words recalled between Trial 5 and Trial 1 from List A; interference = proactive mnemonic interference as measured by total words recalled from List B; immediate recall = total words recalled from List A immediately after interference; delayed recall= total words recalled from List A after the delay. Abbreviations: RSI: restriction spectrum imaging; LRNT: left restricted (intracellular) normalized total fraction, RRNT: right restricted (intracellular) normalized total fraction, LHNT: left hindered (extracellular) normalized total fraction, RHNT: right hindered (extracellular) normalized total fraction, LFNI: left free (water) normalized isotropic fraction, RFNI: right free (water) normalized isotropic fraction, RAVLT: Rey Auditory Verbal Learning Test.

## Discussion

Responding to recent recommendations for source- and composition-specific PM_2.5_ health assessments ^102^, our findings underscore the importance of moving beyond total PM_2.5_ mass to consider specific components and sources, revealing unique links to hippocampal architecture and memory ability during brain development. These findings reflect multivariate covariance patterns, as PLSC identifies coordinated patterns of association across variable sets rather than estimating independent causal mixture effects. Whereas annual average PM_2.5_ total mass was associated with bilateral hippocampal microstructure, including differences in hindered and free water diffusion, component-specific analyses revealed a more nuanced pattern of associations, including between secondary pollutants formed through photochemical reactions (O_3_, SO_4_^2-^), trace elements (Br, V, Fe), and carbons (OC, EC) on hippocampal microstructural properties. PM_2.5_ source analyses further suggest that these associations with hippocampal microstructure may stem from differences in ammonium nitrates, biomass burning, and traffic-related exposures. Moreover, metals (Cu, Zn), largely originating from traffic- and industrial-related emissions, were negatively associated with hippocampal long-axis volumes. We also found converging evidence linking several of these specific pollutants and patterns of hippocampal structure to impaired verbal episodic memory performance. By identifying which components and emission sources are most strongly linked to hippocampal structure and function in pre-adolescents, this work adds to growing evidence that even low levels of air pollution may be harmful to the developing brain ^2,4^.

In terms of toxicity, the health effects of PM_2.5_ are not determined solely by the mass of individual components but by their interactive reactivity ^13,103,104^. Here, we find annual co-exposure, or covariance patterns, of secondary pollutants (O_3_, SO_4_^2-^), trace elements (Br, V, Fe), and both organic and elemental carbons are associated with differences in patterns of intracellular, extracellular, and free water within the hippocampus, suggesting alterations in hippocampal cellularity. Across all hippocampal diffusion analyses, examining PM_2.5_ total mass, components, and sources, a consistent pattern emerged: higher pollution exposure were associated with lower hindered, extracellular diffusion but higher free water diffusion. This pattern may reflect compromised tissue microstructure, as lower external hindered diffusion may indicate a less tortuous, more open extracellular space with fewer cellular obstacles, while higher free water diffusion may suggest increased extracellular fluid in expanded spaces ^55,57,58^. Together, these findings might suggest tissue degradation or atrophy with loss of cellular density, consistent with other known neurodegeneration and/or neuroinflammation processes that occur with air pollution exposure, including changes in neurogenesis, neuron density, gliosis, and/or dendritic branching in response to oxidative stress or neuroinflammation ^2,4,105,106^. In support of this idea, work by Calderón-Garcidueñas and colleagues, which examined children living in highly polluted versus less polluted cities, showed that compared to controls, children with higher exposure exhibited lower left hippocampal NAA/Cr ratios, which they postulate may represent aberrations in hippocampal neuronal density and may be indicative of neuropathology ^107^. Comparatively, rodent models demonstrate that oxidative stress due to O_3_ exposure impacts hippocampal dendritic branching ^108,109^, impacting the cellular environment. Moreover, a recent study using male mice found that exposure to traffic-related UFPs was associated with decreased cell density ^40^ and PM_2.5_ exposure induced reduced neurite density ^39^. Collectively, these findings illustrate how heterogeneous pollutant profiles, even within PM_2.5_ total mass exposure, may be differentially associated with hippocampal tissue characteristics and related neurobiological processes.

Beyond microstructure, we found trace metals, largely originating from industrial and traffic sources, were negatively associated with hippocampal long-axis volumes, with higher annual Cu and Zn exposures being linked to smaller left head and right body and tail volumes. Hippocampal development has been extensively characterized for both gross hippocampal volume ^22–24^ and along its long-axis ^25,27,28^. This developmental heterogeneity suggests that subregions such as the head, body, and tail may differ in their timing of maturation and susceptibility to environmental influences. Here, we saw differential association by hemisphere between hippocampal long-axis volume and annual average Cu and Zn exposure. Moreover, pollutant-specific differences in developmental sensitivity may underlie the spatial patterning of these effects, consistent with evidence that distinct pollutants, such as transition metals, engage different neurotoxic pathways ^37,110^. Given that the hippocampus undergoes protracted change during late childhood and early adolescence, with tradeoffs between head and body volumes ^28,111^, a relatively smaller hippocampal head volume may reflect a delayed maturational trajectory at 9-10-years-old. Recent longitudinal work supports this interpretation, showing that prenatal Cu exposure was associated with smaller total hippocampal volume at age 8 followed by accelerated maturation through age 17 ^33^, suggesting that metal exposures may shift hippocampal developmental timing rather than uniformly impede growth. However, lack of long-axis specific analyses precludes drawing direct comparisons between our results. Evidence suggests metals within PM_2.5_, including Fe, Cu, and Zn can enter the brain through multiple pathways, including direct translocation of UFPs along the olfactory and trigeminal nerves ^15,112^, as well as via systemic circulation via alveolar-capillary transfer in the lungs ^16,112^. These findings may implicate metal dyshomeostasis contributing in part to hippocampal dysfunction as disruptions in metal homeostasis can promote oxidative stress, protein aggregation, and neuroinflammatory cascades that compromise neuronal integrity ^37^.

Beyond its role in driving neuroinflammatory processes ^113^ and its potential for metal dyshomeostasis ^37,114^, air pollution may also affect the brain through dysregulation of the HPA axis ^112,112,115–117^. For example, acute O_3_ exposure has been associated with HPA axis activation and elevated serum levels of corticotropin-releasing hormone, adrenocorticotropic hormone, and cortisol ^118^. Among children and adolescents, O_3_ exposure was also associated with higher serum-derived cortisol levels ^119^. Similarly, inhalation of O_3_ and particulate matter containing polycyclic aromatic hydrocarbons and transition metals increased adrenocorticotropic and glucocorticoid corticosterone levels in animal models, indicating pollutant-induced HPA axis activation ^116^. The hippocampus plays a central role in regulating this stress response system through feedback mechanisms mediated by glucocorticoid receptors, which are densely expressed throughout the hippocampus^120,121^. Under typical conditions, HPA axis activation is transient and promotes adaptation; however, chronic dysregulation may impair hippocampal function and contribute to adverse neurobehavioral outcomes ^122^. Accordingly, PM_2.5_ components and O_3_ may act as toxic stressors that disrupt HPA axis regulation and negatively influence neurocognitive development ^17^. Our observed negative associations between annual O_3_ and OC exposures and differences in extracellular and intracellular diffusion may therefore reflect a consequence of prolonged HPA-axis activation or inflammation-related mechanisms that alter cellular homeostasis within hippocampal tissue. Altogether, these findings highlight the pervasive influence of air pollution on the hippocampus and raise questions regarding implications for developmental and lifespan hippocampal integrity.

We included the RAVLT to assess whether the structural associations observed between air pollution and the hippocampus extended to hippocampal-dependent memory performance. Epidemiological and experimental work reinforce this idea, with recent research demonstrating that air pollution adversely affects cognitive function throughout the lifespan ^6,7,9^, and rodent studies showing that exposure to UFPs and PM_2.5_ disrupts hippocampal-dependent learning and memory ^38,39^. Here, we found converging evidence linking specific pollutants to both hippocampal structure and cognition. For example, higher Cu and Zn from traffic and industrial-related sources, was associated with smaller hippocampal volumes, and higher Zn, in particular, was linked to poorer learning and immediate recall performance.

We also found smaller hippocampal tail volumes were associated with poorer immediate recall, suggesting that structural alterations within the hippocampus may have functional consequences for verbal episodic memory. Immediate recall performance is thought to reflect encoding efficiency and early consolidation, relying on hippocampal function, whereas delayed recall additionally reflects retrieval and distributed cortical-hippocampal network interactions that may be less directly captured by the hippocampus alone ^21,26,34^. Immediate recall may thus be more sensitive to subtle neurodevelopmental differences than delayed recall. Notably, hippocampal tail regions showed the strongest associations with memory performance. Posterior hippocampal regions, including the tail, support detailed episodic representations and undergo protracted structural refinement during late childhood and early adolescence ^20,25–28^. Consistent with a developmental framework, prior work has shown that larger hippocampal tail volume is associated with better episodic memory performance in children but not adults, suggesting that posterior hippocampal regions may play a particularly important role in supporting memory during this developmental period ^20^. An RSI study investigating cognitive impairment lend further support for this interpretation, reporting that reduced neurite density (lower RNT) and increased free water (higher FNI) in white matter tracts projecting to the hippocampus were associated with poorer memory, reinforcing the functional relevance of hippocampal connectivity to episodic learning ^123^. In our current study, we found that both PM_2.5_ total mass, as well as several components, were linked to higher free water alongside lower hindered, extracellular signal fractions within the hippocampus. In turn, this pattern of microstructure in the hippocampus was associated with poorer immediate and delayed recall performance. Together these findings highlight possible pollutant-specific patterns of covariance between brain and behavior. While our PLSC models capture these complex multivariate relationships (i.e., between air quality and brain, air quality and behavior, and brain and behavior), PLSC does not allow for testing mediation directly; thus, future research is needed to determine whether hippocampal architecture statistically mediates the relationship between air pollution and cognitive performance.

Moving forward, understanding how air pollution impacts cognition will require careful attention to the types of pollutants involved and the specific cognitive processes assessed. In highly air pollution-exposed urban youth, air pollution has been linked to impaired attention, short-term memory, and learning performance ^49^. Additionally, findings from the BREATHE cohort showed that exposure to black carbon and PM_2.5_ predicted declines in working memory for 7-10-year-olds over a 12-month period ^50^, and a recent meta-analysis confirmed that greater PM_2.5_ total mass exposure is associated with poorer working memory performance in childhood ^7^. Moreover, prior work from our group using a mixture modeling approach, known as weighted quantile sum, found that ammonium nitrates were associated with reduced performance on a composite measure of working and episodic learning and memory, whereas traffic-related pollutants were associated with poorer executive functioning ^10^. In this previous work, we utilized composite measures of cognition, created using a principal component analysis of the ABCD cognitive battery ^124^, which included list learning from the NIH toolbox as well as only the total number of words learned from the RAVLT task, whereas the executive function domain included NIH toolbox tasks of Flanker, Card-Sorting, and Pattern Comparison ^124^. Because studies assess different cognitive abilities at different ages using distinct tasks and analytic approaches, the emerging literature shows that while air pollution is related to cognitive ability, the specific pattern of findings depends on what is measured, how it is measured, and which pollutants are considered.

Altogether, in combination with the extant literature our findings suggest a broader vulnerability of hippocampal-dependent function to air pollution exposure in childhood that may ‘set the stage’ for later-life memory ability. In humans, the detrimental effects of air pollution on the hippocampus and declines in cognitive function during adulthood are increasingly recognized ^7,9,125^. Among older adults, single-pollutant models of PM_2.5_ total mass, SO_2_, NO_2_, and O_3_ were independently associated with poorer list learning and memory ability ^126^. Given the hippocampus’s role in memory ability and its fixture in neurodegenerative diseases ^127,128^, it is crucial to take a lifespan framework approach to investigating early-life factors that may contribute to neurocognitive health. Among Mexico City highly exposed children and adolescents compared to less-exposed controls, higher levels of air pollution exposure was linked to Alzheimer’s disease-related neuropathology and impairments in cognitive performance ^114,129,130^, indicating pathology typically seen in older adulthood may exist as early as adolescence. A recent meta-analysis identified PM_2.5_ as an environmental determinant of dementia including Alzheimer’s disease^131^ and linked pollution exposure to cognitive decline^132^. Additionally, disruptions in the homeostasis of essential transition metals (e.g., Cu, Zn) have been implicated in neurodegenerative diseases such as Alzheimer’s and Parkinson’s ^37^, with copper dyshomeostasis, in particular, possibly playing a central role ^110^. Mirroring our current findings on source-specific exposure, recent research in older adulthood has shown that biomass burning and ammonium nitrates show the strongest links with incident dementia ^133^. Thus, studying how air pollution may impact hippocampal architecture and cognition in childhood may present a key opportunity to focus public health strategies toward the prevention of neurocognitive impairments throughout adolescence into adulthood ^18^, supporting the success of children as they develop.

### Strengths & Limitations

Our approach provides a comprehensive view of exposure to multiple pollutants and hippocampal architecture by investigating hippocampal micro- and macro-structure. This nuanced analysis offers valuable insights into the impact of air pollution on hippocampal structure during the sensitive period of pre-adolescence. However, some limitations warrant consideration. First, analyses were constrained to the cross-sectional measures available in the public ABCD Study dataset, limiting our ability to investigate lifetime air pollution history and additional measures of hippocampal structure and function. Second, we could not examine diffusion properties within specific hippocampal long-axis subregions which would have allowed us to disentangle specific differences in cell type or presence of white matter (e.g., alveus located on the hippocampal body). However, emerging neuroimaging techniques ^134^ will enable future studies to explore these finer-scale distinctions. Third, the cross-sectional design limits our ability to draw causal inferences and track developmental trajectories. Although we were able to show that relationships exist between air pollution, hippocampal architecture, and memory ability, future studies that employ longitudinal or multivariate mediation analyses are needed to disentangle the complex pathways linking air pollution exposure to cognition. Moreover, studies that examine lifetime exposures could provide more detailed information as to which periods of air pollution exposure during childhood might shape hippocampal structure and dependent behavior. Fourth, the ABCD Study does not include measures of indoor nor school air pollution exposure. However, PM_2.5_ predominantly contributes to indoor particulate matter concentrations (outdoor-to-indoor ratios: 0.5-0.8 for PM_2.5_)^135,136^. Nonetheless, indoor sources of air pollution represent an additional exposure pathway not captured in the present study and may introduce some degree of exposure misclassification. Fifth, sex-specific differences in air pollution-brain associations were not examined in the present study and represent an important direction for future research. Finally, although RSI has been histologically validated in several subcortical structures containing mixed cellular and myelinated tissue ^57^, similar validation studies have not yet been conducted in the hippocampus. This absence of direct histological correspondence limits our ability to determine the cellular or cytoarchitectural basis of hippocampal diffusion measures, underscoring the need for future multimodal and postmortem work to establish the biological ground truth of hippocampal microstructure using this method.

## Conclusions

This study adds to the literature by showing that the link between air pollution and hippocampal structure is not only related to PM_2.5_ total mass levels, which are commonly used in research and regulation, but is also driven by component and source-specific exposures. Particularly, PM_2.5_ constituents and their related sources are more closely linked to differences in hippocampal structure and verbal episodic memory performance as compared to total PM_2.5_ mass, highlighting both structural and functional sensitivity to ambient air quality. By identifying which components and emission sources are most strongly linked to neurodevelopmental differences, this work adds to growing evidence that even low levels of air pollution can influence brain structure and function. Additional studies examining co-exposure to PM_2.5_ components and sources are essential to pave a clearer path toward targeted and effective public health action to reduce harmful exposures and promote cognitive health across childhood and adolescence.

## Acknowledgments

A special thank you to all participants and their families for their participation in the ABCD Study.

## Data Availability

Data used in the preparation of this article were obtained from the Adolescent Brain Cognitive DevelopmentSM (ABCD) Study (https://abcdstudy.org), held in the NIMH Data Archive (NDA). This is a multisite, longitudinal study designed to recruit more than 10,000 children aged 9-10 and follow them over 10 years into early adulthood. The ABCD Study® is supported by the National Institutes of Health and additional federal partners under award numbers U01DA041048, U01DA050989, U01DA051016, U01DA041022, U01DA051018, U01DA051037, U01DA050987, U01DA041174, U01DA041106, U01DA041117, U01DA041028, U01DA041134, U01DA050988, U01DA051039, U01DA041156, U01DA041025, U01DA041120, U01DA051038, U01DA041148, U01DA041093, U01DA041089, U24DA041123, U24DA041147. A full list of supporters is available at https://abcdstudy.org/federal-partners.html. A listing of participating sites and a complete listing of the study investigators can be found at https://abcdstudy.org/consortium_members/. ABCD consortium investigators designed and implemented the study and/or provided data but did not necessarily participate in the analysis or writing of this report. This manuscript reflects the views of the authors and may not reflect the opinions or views of the NIH or ABCD consortium investigators.

The ABCD data repository grows and changes over time. The ABCD data used in this report came from [doi:10.15154/8873-zj65]. DOIs can be found at [https://nda.nih.gov/study.html?id=2147].

## Author Contributions

Michael A. Rosario: Formal Analysis, Writing - Original Draft and Editing, Visualization. Kirthana Sukumaran: Writing - Review & Editing, Statistical Supervision. Kathryn Bottenhorn: Writing - Review & Editing. Alethea de Jesus: Data Curation, Writing - Review & Editing. Hedyeh Ahmadi: Statistical Supervision, Writing - Review & Editing. Carlos Cardenas-Inigues: Writing - Review & Editing, Statistical Supervision. Rima Habre: Methodology, Writing - Review & Editing. Shermaine Abad: Data Curation, Writing – Review & Editing. Jacob G. Pine: Data Curation, Writing - Review & Editing. Deanna M. Barch: Data Curation, Writing - Review & Editing. Joel Schwartz: Methodology, Data Curation, Resources, Writing - Review & Editing. Daniel A. Hackman: Methodology, Writing – Review & Editing. Jiu-Chiuan Chen: Conceptualization, Methodology, Writing - Review & Editing. Megan M. Herting: Funding Acquisition, Conceptualization, Methodology, Supervision, Project Administration, Writing - Review & Editing.

## References

1 Brumberg HL, Karr CJ, Bole A, et al. Ambient Air Pollution: Health Hazards to Children. Pediatrics 2021; 147: e2021051484.

2 Cory-Slechta DA, Merrill A, Sobolewski M. Air Pollution–Related Neurotoxicity Across the Life Span. Annu Rev Pharmacol Toxicol 2023; 63: 143–63.

3 Fuller R, Landrigan PJ, Balakrishnan K, et al. Pollution and health: a progress update. Lancet Planet Health 2022; 6: e535–47.

4 Herting MM, Bottenhorn KL, Cotter DL. Outdoor air pollution and brain development in childhood and adolescence. Trends Neurosci 2024; 47: 593–607.

5 Morrel J, Dong M, Rosario MA, Cotter DL, Bottenhorn KL, Herting MM. A Systematic Review of Air Pollution Exposure and Brain Structure and Function during Development. Environ Res 2025; : 121368.

6 Alter NC, Whitman EM, Bellinger DC, Landrigan PJ. Quantifying the association between PM2.5 air pollution and IQ loss in children: a systematic review and meta-analysis. Environ Health 2024; 23: 101.

7 Thompson R, Smith RB, Karim YB, et al. Air pollution and human cognition: A systematic review and meta-analysis. Sci Total Environ 2023; 859: 160234.

8 US EPA O. Criteria Air Pollutants. 2014; published online April 9. https://www.epa.gov/criteria-air-pollutants (accessed Feb 4, 2025).

9 Geto AK, Feleke SF, Yimer A, et al. The association between air pollution and cognitive impairment: a systematic review and meta-analysis of global studies. BMC Public Health 2025; 25: 3548.

10 Sukumaran K, Botternhorn KL, Schwartz J, et al. Associations between Fine Particulate Matter Components, Their Sources, and Cognitive Outcomes in Children Ages 9–10 Years Old from the United States. Environ Health Perspect 2024; 132: 107009.

11 US EPA O. National Ambient Air Quality Standards (NAAQS) for PM. 2020; published online April 13. https://www.epa.gov/pm-pollution/national-ambient-air-quality-standards-naaqs-pm (accessed Nov 20, 2024).

12 Europe WHORO for. Health relevance of particulate matter from various sources : report on a WHO workshop, Bonn, Germany 26-27 March 2007. 2007. https://iris.who.int/handle/10665/107846 (accessed July 15, 2024).

13 Shang J, Zhang Y, Schauer JJ, et al. Prediction of the oxidation potential of PM2.5 exposures from pollutant composition and sources. Environ Pollut 2022; 293: 118492.

14 Park M, Joo HS, Lee K, et al. Differential toxicities of fine particulate matters from various sources. Sci Rep 2018; 8: 17007.

15 Block ML, Calderón-Garcidueñas L. Air pollution: mechanisms of neuroinflammation and CNS disease. Trends Neurosci 2009; 32: 506–16.

16 Ku T, Zhang Y, Ji X, Li G, Sang N. PM2.5-bound metal metabolic distribution and coupled lipid abnormality at different developmental windows. Environ Pollut 2017; 228: 354–62.

17 Thomson EM. Neurobehavioral and metabolic impacts of inhaled pollutants: A role for the hypothalamic-pituitary-adrenal axis? Endocr Disruptors 2013; 1: e27489.

18 Farina FR, Bridgeman K, Gregory S, et al. Next generation brain health: transforming global research and public health to promote prevention of dementia and reduce its risk in young adult populations. Lancet Healthy Longev 2024; 5: 100665.

19 Botdorf M, Canada KL, Riggins T. A meta-analysis of the relation between hippocampal volume and memory ability in typically developing children and adolescents. Hippocampus 2022; 32: 386–400.

20 DeMaster D, Pathman T, Lee JK, Ghetti S. Structural development of the hippocampus and episodic memory: developmental differences along the anterior/posterior axis. Cereb Cortex N Y N 1991 2014; 24: 3036–45.

21 Dickerson BC, Eichenbaum H. The episodic memory system: neurocircuitry and disorders. Neuropsychopharmacol Off Publ Am Coll Neuropsychopharmacol 2010; 35: 86–104.

22 Gogtay N, Nugent III TF, Herman DH, et al. Dynamic mapping of normal human hippocampal development. Hippocampus 2006; 16: 664–72.

23 Herting MM, Johnson C, Mills KL, et al. Development of subcortical volumes across adolescence in males and females: A multisample study of longitudinal changes. NeuroImage 2018; 172: 194–205.

24 Uematsu A, Matsui M, Tanaka C, et al. Developmental Trajectories of Amygdala and Hippocampus from Infancy to Early Adulthood in Healthy Individuals. PLoS ONE 2012; 7: e46970.

25 Dick AS, Ralph Y, Farrant K, Reeb-Sutherland B, Pruden S, Mattfeld A. Volumetric development of hippocampal subfields and hippocampal white matter connectivity: Relationship with episodic memory. Dev Psychobiol 2022; 64: e22333.

26 Poppenk J, Evensmoen HR, Moscovitch M, Nadel L. Long-axis specialization of the human hippocampus. Trends Cogn Sci 2013; 17: 230–40.

27 Riggins T, Blankenship SL, Mulligan E, Rice K, Redcay E. Developmental differences in relations between episodic memory and hippocampal subregion volume during early childhood. Child Dev 2015; 86: 1710–8.

28 Tamnes CK, Bos MGN, van de Kamp FC, Peters S, Crone EA. Longitudinal development of hippocampal subregions from childhood to adulthood. Dev Cogn Neurosci 2018; 30: 212–22.

29 Binter A-C, Kusters MSW, van den Dries MA, et al. Air pollution, white matter microstructure, and brain volumes: Periods of susceptibility from pregnancy to preadolescence. Environ Pollut Barking Essex 1987 2022; 313: 120109.

30 Buthmann JL, Benmarhnia T, Huang JY, et al. Exposure to Fine Particulate Matter During Pregnancy is Associated with Hippocampal Development in Offspring. Biol Psychiatry Glob Open Sci 2025; : 100490.

31 Cserbik D, Chen J-C, McConnell R, et al. Fine particulate matter exposure during childhood relates to hemispheric-specific differences in brain structure. Environ Int 2020; 143: 105933.

32 Sukumaran K, Bottenhorn KL, Rosario MA, et al. Sources and components of fine air pollution exposure and brain morphology in preadolescents. Sci Total Environ 2025; 979: 179448.

33 Kusters MSW, Binter A-C, Muetzel RL, et al. Outdoor residential air pollution exposure and the development of brain volumes across childhood: A longitudinal study. Environ Pollut 2025; 373: 126078.

34 Knierim JJ. The hippocampus. Curr Biol 2015; 25: R1116–21.

35 Karat BG, DeKraker J, Hussain U, Köhler S, Khan AR. Mapping the macrostructure and microstructure of the in vivo human hippocampus using diffusion MRI. Hum Brain Mapp 2023; 44: 5485–503.

36 Karat BG, Genc S, Raven EP, Palombo M, Khan AR, Jones DK. The developing hippocampus: Microstructural evolution through childhood and adolescence. BioRxiv Prepr Serv Biol 2024; : 2024.08.19.608590.

37 Cory-Slechta DA, Sobolewski M, Oberdörster G. Air Pollution-Related Brain Metal Dyshomeostasis as a Potential Risk Factor for Neurodevelopmental Disorders and Neurodegenerative Diseases. Atmosphere 2020; 11: 1098.

38 Fonken LK, Xu X, Weil ZM, et al. Air pollution impairs cognition, provokes depressive-like behaviors and alters hippocampal cytokine expression and morphology. Mol Psychiatry 2011; 16: 987–95.

39 Woodward NC, Haghani A, Johnson RG, et al. Prenatal and early life exposure to air pollution induced hippocampal vascular leakage and impaired neurogenesis in association with behavioral deficits. Transl Psychiatry 2018; 8: 1–10.

40 Ehsanifar M, Yavari Z, Rafati M. Exposure to urban air pollution particulate matter: neurobehavioral alteration and hippocampal inflammation. Environ Sci Pollut Res 2022; 29: 50856–66.

41 Patten KT, González EA, Valenzuela A, et al. Effects of early life exposure to traffic-related air pollution on brain development in juvenile Sprague-Dawley rats. Transl Psychiatry 2020; 10: 166.

42 Berg EL, Pedersen LR, Pride MC, et al. Developmental exposure to near roadway pollution produces behavioral phenotypes relevant to neurodevelopmental disorders in juvenile rats. Transl Psychiatry 2020; 10: 289.

43 Bos B, Barratt B, Batalle D, et al. Prenatal exposure to air pollution is associated with structural changes in the neonatal brain. Environ Int 2023; 174: 107921.

44 Essers E, Binter A-C, Neumann A, White T, Alemany S, Guxens M. Air pollution exposure during pregnancy and childhood, APOE ε4 status and Alzheimer polygenic risk score, and brain structural morphology in preadolescents. Environ Res 2023; 216: 114595.

45 Lubczyńska MJ, Muetzel RL, El Marroun H, et al. Air pollution exposure during pregnancy and childhood and brain morphology in preadolescents. Environ Res 2021; 198: 110446.

46 Herting MM, Burnor E, Ahmadi H, et al. Air Pollution Exposure, Prefrontal Connectivity, and Emotional Behavior in Early Adolescence. Res Rep Health Eff Inst 2025; 2025: 225.

47 Hamra GB, Buckley JP. Environmental Exposure Mixtures: Questions and Methods to Address Them. Curr Epidemiol Rep 2018; 5: 160–5.

48 Burnor E, Cserbik D, Cotter DL, et al. Association of Outdoor Ambient Fine Particulate Matter With Intracellular White Matter Microstructural Properties Among Children. JAMA Netw Open 2021; 4: e2138300.

49 Calderón-Garcidueñas L, Engle R, Mora-Tiscareño A, et al. Exposure to severe urban air pollution influences cognitive outcomes, brain volume and systemic inflammation in clinically healthy children. Brain Cogn 2011; 77: 345–55.

50 Alvarez-Pedrerol M, Rivas I, López-Vicente M, et al. Impact of commuting exposure to traffic-related air pollution on cognitive development in children walking to school. Environ Pollut 2017; 231: 837–44.

51 Garavan H, Bartsch H, Conway K, et al. Recruiting the ABCD sample: Design considerations and procedures. Dev Cogn Neurosci 2018; 32: 16–22.

52 Volkow ND, Koob GF, Croyle RT, et al. The conception of the ABCD study: From substance use to a broad NIH collaboration. Dev Cogn Neurosci 2018; 32: 4–7.

53 Bottenhorn KL, Sukumaran K, Cardenas-Iniguez C, et al. Air pollution from biomass burning disrupts early adolescent cortical microarchitecture development. Environ Int 2024; 189: 108769.

54 Jin T, Amini H, Kosheleva A, et al. Associations between long-term exposures to airborne PM2.5 components and mortality in Massachusetts: mixture analysis exploration. Environ Health 2022; 21: 96.

55 Palmer CE, Pecheva D, Iversen JR, et al. Microstructural development from 9 to 14 years: Evidence from the ABCD Study. Dev Cogn Neurosci 2021; 53: 101044.

56 Pecheva D, Smith DM, Casey BJ, et al. Sex and mental health are related to subcortical brain microstructure. Proc Natl Acad Sci U S A 2024; 121: e2403212121.

57 White NS, Leergaard TB, D’Arceuil H, Bjaalie JG, Dale AM. Probing tissue microstructure with restriction spectrum imaging: Histological and theoretical validation. Hum Brain Mapp 2013; 34: 327–46.

58 White NS, McDonald CR, Farid N, et al. Diffusion-Weighted Imaging in Cancer: Physical Foundations and Applications of Restriction Spectrum Imaging. Cancer Res 2014; 74: 4638–52.

59 Fischl B, Salat DH, Busa E, et al. Whole Brain Segmentation: Automated Labeling of Neuroanatomical Structures in the Human Brain. Neuron 2002; 33: 341–55.

60 Iglesias JE, Augustinack JC, Nguyen K, et al. A computational atlas of the hippocampal formation using ex vivo, ultra-high resolution MRI: Application to adaptive segmentation of in vivo MRI. NeuroImage 2015; 115: 117–37.

61 Pine JG, Paul SE, Johnson E, Bogdan R, Kandala S, Barch DM. Polygenic Risk for Schizophrenia, Major Depression, and Post-traumatic Stress Disorder and Hippocampal Subregion Volumes in Middle Childhood. Behav Genet 2023; 53: 279–91.

62 Gard AM, Hyde LW, Heeringa SG, West BT, Mitchell C. Why weight? Analytic approaches for large-scale population neuroscience data. Dev Cogn Neurosci 2023; 59: 101196.

63 Fan CC, Marshall A, Smolker H, et al. Adolescent Brain Cognitive Development (ABCD) study Linked External Data (LED): Protocol and practices for geocoding and assignment of environmental data. Dev Cogn Neurosci 2021; 52: 101030.

64 Di Q, Amini H, Shi L, et al. An ensemble-based model of PM2.5 concentration across the contiguous United States with high spatiotemporal resolution. Environ Int 2019; 130: 104909.

65 Di Q, Amini H, Shi L, et al. Assessing NO2 Concentration and Model Uncertainty with High Spatiotemporal Resolution across the Contiguous United States Using Ensemble Model Averaging. Environ Sci Technol 2020; 54: 1372–84.

66 Requia WJ, Di Q, Silvern R, et al. An Ensemble Learning Approach for Estimating High Spatiotemporal Resolution of Ground-Level Ozone in the Contiguous United States. Environ Sci Technol 2020; 54: 11037–47.

67 Amini H, Danesh-Yazdi M, Di Q, et al. Hyperlocal super-learned PM2.5 components across the contiguous US. 2022; published online Dec 28. DOI:10.21203/rs.3.rs-1745433/v2.

68 Amini H, Danesh-Yazdi M, Di Q, et al. Annual Mean PM2. 5 Components (EC, NH4, NO3, OC, SO4) 50 m Urban and 1 km Non-Urban Area Grids for Contiguous US, 2000-2019 v1.(Preliminary Release). NASA Socioeconomic Data and Applications Center (SEDAC) Palisades, NY, 2022.

69 Amini H, Danesh-Yazdi M, Di Q, et al. Annual Mean PM2.5 Trace Elements 50m Grids in Urban Areas and 1km Grids in Non-Urban Areas for Contiguous U.S., 2000-2019, v1. 2023. DOI:10.7927/1X94-MV38.

70 Feng Y, Castro E, Wei Y, et al. Long-term exposure to ambient PM2.5, particulate constituents and hospital admissions from non-respiratory infection. Nat Commun 2024; 15: 1518.

71 Wang Y, Shi L, Lee M, et al. Long-term Exposure to PM2.5 and Mortality Among Older Adults in the Southeastern US. Epidemiology 2017; 28: 207.

72 Cory-Slechta DA, Marvin E, Welle K, Oberdörster G, Sobolewski M. The protracted neurotoxic consequences in mice of developmental exposures to inhaled iron nanoparticles alone or in combination with SO2. Front Behav Neurosci 2025; 19. DOI:10.3389/fnbeh.2025.1544974.

73 Norris G, Duvall R, Brown S, Bai S. EPA positive matrix factorization (PMF) 5.0 fundamentals and user guide, US Environmental Protection Agency, Washington, DC. Www2 Epa Govsitesproductionfiles2015-02documentspmf5 0userguide Pdf 2014.

74 Casey BJ, Cannonier T, Conley MI, et al. The Adolescent Brain Cognitive Development (ABCD) study: Imaging acquisition across 21 sites. Dev Cogn Neurosci 2018; 32: 43–54.

75 Hagler DJ, Hatton S, Cornejo MD, et al. Image processing and analysis methods for the Adolescent Brain Cognitive Development Study. NeuroImage 2019; 202: 116091.

76 Moeller S, Yacoub E, Olman CA, et al. Multiband multislice GE-EPI at 7 tesla, with 16-fold acceleration using partial parallel imaging with application to high spatial and temporal whole-brain fMRI. Magn Reson Med 2010; 63: 1144–53.

77 Setsompop K, Cohen-Adad J, Gagoski BA, et al. Improving diffusion MRI using simultaneous multi-slice echo planar imaging. NeuroImage 2012; 63: 569–80.

78 Calderón-Garcidueñas L, Hernández-Luna J, Mukherjee PS, et al. Hemispheric Cortical, Cerebellar and Caudate Atrophy Associated to Cognitive Impairment in Metropolitan Mexico City Young Adults Exposed to Fine Particulate Matter Air Pollution. Toxics 2022; 10: 156.

79 Peterson BS, Rauh VA, Bansal R, et al. Effects of Prenatal Exposure to Air Pollutants (Polycyclic Aromatic Hydrocarbons) on the Development of Brain White Matter, Cognition, and Behavior in Later Childhood. JAMA Psychiatry 2015; 72: 531–40.

80 Lampinen B, Szczepankiewicz F, Mårtensson J, van Westen D, Sundgren PC, Nilsson M. Neurite density imaging versus imaging of microscopic anisotropy in diffusion MRI: A model comparison using spherical tensor encoding. NeuroImage 2017; 147: 517–31.

81 Zhang H, Schneider T, Wheeler-Kingshott CA, Alexander DC. NODDI: practical in vivo neurite orientation dispersion and density imaging of the human brain. NeuroImage 2012; 61: 1000–16.

82 Cotter DL, Ahmadi H, Cardenas-Iniguez C, et al. Sex-specific effects in how childhood exposures to multiple ambient air pollutants affect white matter microstructure development across early adolescence. 2023; published online Aug 17. DOI:10.21203/rs.3.rs-3213618/v1.

83 Cotter DL, Ahmadi H, Cardenas-Iniguez C, et al. Exposure to multiple ambient air pollutants changes white matter microstructure during early adolescence with sex-specific differences. Commun Med 2024; 4: 1–12.

84 Sukumaran K, Cardenas-Iniguez C, Burnor E, et al. Ambient fine particulate exposure and subcortical gray matter microarchitecture in 9- and 10-year-old children across the United States. iScience 2023; 26: 106087.

85 Conley MI, Skalaban LJ, Rapuano KM, et al. Altered hippocampal microstructure and function in children who experienced Hurricane Irma. Dev Psychobiol 2021; 63: 864–77.

86 Li ZA, Cai Y, Taylor RL, et al. Associations Between Socioeconomic Status, Obesity, Cognition, and White Matter Microstructure in Children. JAMA Netw Open 2023; 6: e2320276.

87 Luciana M, Bjork JM, Nagel BJ, et al. Adolescent neurocognitive development and impacts of substance use: Overview of the adolescent brain cognitive development (ABCD) baseline neurocognition battery. Dev Cogn Neurosci 2018; 32: 67–79.

88 Peaker A, Stewart LE. Rey’s Auditory Verbal Learning Test — A Review. In: Crawford JR, Parker DM, eds. Developments in Clinical and Experimental Neuropsychology. Boston, MA: Springer US, 1989: 219–36.

89 Beaton D, Chin Fatt CR, Abdi H. An ExPosition of multivariate analysis with the singular value decomposition in R. Comput Stat Data Anal 2014; 72: 176–89.

90 Abdi H, Williams LJ. Partial Least Squares Methods: Partial Least Squares Correlation and Partial Least Square Regression. In: Computational Toxicology. Humana Press, Totowa, NJ, 2013: 549–79.

91 Krishnan A, Williams LJ, McIntosh AR, Abdi H. Partial Least Squares (PLS) methods for neuroimaging: a tutorial and review. NeuroImage 2011; 56: 455–75.

92 Hajat A, Hsia C, O’Neill MS. Socioeconomic Disparities and Air Pollution Exposure: a Global Review. Curr Environ Health Rep 2015; 2: 440–50.

93 Nunez Y, Benavides J, Shearston JA, et al. An environmental justice analysis of air pollution emissions in the United States from 1970 to 2010. Nat Commun 2024; 15: 268.

94 US Census Bureau. 2010 Urban Area FAQs. Census.gov. https://www.census.gov/programs-surveys/geography/about/faq/2010-urban-area-faq.html (accessed Nov 23, 2025).

95 Echeverria SE, Diez-Roux AV, Link BG. Reliability of self-reported neighborhood characteristics. J Urban Health Bull N Y Acad Med 2004; 81: 682–701.

96 McIntosh AR, Lobaugh NJ. Partial least squares analysis of neuroimaging data: applications and advances. NeuroImage 2004; 23: S250–63.

97 Abdi H, Beaton D. Principal Component and Correspondence Analyses Using R. Springer, 2015 https://link.springer.com/book/9783319092553 (accessed Nov 23, 2025).

98 Griffis JC, Metcalf NV, Corbetta M, Shulman GL. Structural Disconnections Explain Brain Network Dysfunction after Stroke. Cell Rep 2019; 28: 2527–2540.e9.

99 Mišić B, Betzel RF, de Reus MA, et al. Network-Level Structure-Function Relationships in Human Neocortex. Cereb Cortex N Y N 1991 2016; 26: 3285–96.

100 US EPA O. NAAQS Table. 2014; published online April 10. https://www.epa.gov/criteria-air-pollutants/naaqs-table (accessed Nov 23, 2025).

101 World Health Organization. What are the WHO Air quality guidelines? World Health Organ. 2021; published online Sept 22. https://www.who.int/news-room/feature-stories/detail/what-are-the-who-air-quality-guidelines (accessed Jan 29, 2025).

102 Vilcassim R, Thurston GD. Gaps and future directions in research on health effects of air pollution. eBioMedicine 2023; 93: 104668.

103 Kelly FJ, Fussell JC. Size, source and chemical composition as determinants of toxicity attributable to ambient particulate matter. Atmos Environ 2012; 60: 504–26.

104 Lin Y-C, Li Y-C, Amesho KTT, Shangdiar S, Chou F-C, Cheng P-C. Chemical characterization of PM2.5 emissions and atmospheric metallic element concentrations in PM2.5 emitted from mobile source gasoline-fueled vehicles. Sci Total Environ 2020; 739: 139942.

105 Kang YJ, Tan H-Y, Lee CY, Cho H. An Air Particulate Pollutant Induces Neuroinflammation and Neurodegeneration in Human Brain Models. Adv Sci 2021; 8: 2101251.

106 Liu F, Liu C, Liu Y, Wang J, Wang Y, Yan B. Neurotoxicity of the air-borne particles: From molecular events to human diseases. J Hazard Mater 2023; 457: 131827.

107 Calderón-Garcidueñas L, Mora-Tiscareño A, Melo-Sánchez G, et al. A Critical Proton MR Spectroscopy Marker of Alzheimer’s Disease Early Neurodegenerative Change: Low Hippocampal NAA/Cr Ratio Impacts APOE ɛ4 Mexico City Children and Their Parents. J Alzheimer’s Dis 2015; 48: 1065–75.

108 Bello-Medina PC, Prado-Alcalá RA, Rivas-Arancibia S. Effect of Ozone Exposure on Dendritic Spines of CA1 Pyramidal Neurons of the Dorsal Hippocampus and on Object–place Recognition Memory in Rats. Neuroscience 2019; 402: 1–10.

109 Rivas-Arancibia S, Guevara-Guzmán R, López-Vidal Y, et al. Oxidative Stress Caused by Ozone Exposure Induces Loss of Brain Repair in the Hippocampus of Adult Rats. Toxicol Sci 2010; 113: 187–97.

110 Manto M. Abnormal Copper Homeostasis: Mechanisms and Roles in Neurodegeneration. Toxics 2014; 2: 327–45.

111 Riggins T, Geng F, Botdorf M, Canada K, Cox L, Hancock GR. Protracted hippocampal development is associated with age-related improvements in memory during early childhood. NeuroImage 2018; 174: 127–37.

112 Heusinkveld HJ, Wahle T, Campbell A, et al. Neurodegenerative and neurological disorders by small inhaled particles. NeuroToxicology 2016; 56: 94–106.

113 Brockmeyer S, D’Angiulli A. How air pollution alters brain development: the role of neuroinflammation. Transl Neurosci 2016; 7: 24–30.

114 Calderón-Garcidueñas L, Serrano-Sierra A, Torres-Jardón R, et al. The impact of environmental metals in young urbanites’ brains. Exp Toxicol Pathol 2013; 65: 503–11.

115 McEwen BS, Tucker P. Critical Biological Pathways for Chronic Psychosocial Stress and Research Opportunities to Advance the Consideration of Stress in Chemical Risk Assessment. Am J Public Health 2011; 101: S131–9.

116 Thomson EM, Vladisavljevic D, Mohottalage S, Kumarathasan P, Vincent R. Mapping Acute Systemic Effects of Inhaled Particulate Matter and Ozone: Multiorgan Gene Expression and Glucocorticoid Activity. Toxicol Sci 2013; 135: 169–81.

117 Thomson EM, Pal S, Guénette J, et al. Ozone Inhalation Provokes Glucocorticoid-Dependent and -Independent Effects on Inflammatory and Metabolic Pathways. Toxicol Sci 2016; 152: 17–28.

118 Xia Y, Niu Y, Cai J, et al. Personal ozone exposure and stress hormones in the hypothalamus–pituitary–adrenal and sympathetic-adrenal-medullary axes. Environ Int 2022; 159: 107050.

119 Toledo-Corral CM, Alderete TL, Herting MM, et al. Ambient air pollutants are associated with morning serum cortisol in overweight and obese Latino youth in Los Angeles. Environ Health 2021; 20: 39.

120 Seckl JR, Dickson KL, Yates C, Fink G. Distribution of glucocorticoid and mineralocorticoid receptor messenger RNA expression in human postmortem hippocampus. Brain Res 1991; 561: 332–7.

121 Wang Q, Van Heerikhuize J, Aronica E, et al. Glucocorticoid receptor protein expression in human hippocampus; stability with age. Neurobiol Aging 2013; 34: 1662–73.

122 McEwen BS, Nasca C, Gray JD. Stress Effects on Neuronal Structure: Hippocampus, Amygdala, and Prefrontal Cortex. Neuropsychopharmacology 2016; 41: 3–23.

123 Reas ET, Hagler DJ, White NS, et al. Sensitivity of restriction spectrum imaging to memory and neuropathology in Alzheimer’s disease. Alzheimers Res Ther 2017; 9: 55.

124 Thompson WK, Barch DM, Bjork JM, et al. The structure of cognition in 9 and 10 year-old children and associations with problem behaviors: Findings from the ABCD study’s baseline neurocognitive battery. Dev Cogn Neurosci 2019; 36: 100606.

125 Hedges DW, Erickson LD, Kunzelman J, Brown BL, Gale SD. Association between exposure to air pollution and hippocampal volume in adults in the UK Biobank. NeuroToxicology 2019; 74: 108–20.

126 Shin J, Han S-H, Choi J, Shin J, Han S-H, Choi J. Exposure to Ambient Air Pollution and Cognitive Impairment in Community-Dwelling Older Adults: The Korean Frailty and Aging Cohort Study. Int J Environ Res Public Health 2019; 16. DOI:10.3390/ijerph16193767.

127 Dugger BN, Dickson DW. Pathology of Neurodegenerative Diseases. Cold Spring Harb Perspect Biol 2017; 9: a028035.

128 Small SA, Schobel SA, Buxton RB, Witter MP, Barnes CA. A pathophysiological framework of hippocampal dysfunction in ageing and disease. Nat Rev Neurosci 2011; 12: 585–601.

129 Calderón-Garcidueñas L, Mora-Tiscareño A, Ontiveros E, et al. Air pollution, cognitive deficits and brain abnormalities: A pilot study with children and dogs. Brain Cogn 2008; 68: 117–27.

130 Calderón-Garcidueñas L, González-Maciel A, Reynoso-Robles R, et al. Alzheimer’s disease and alpha-synuclein pathology in the olfactory bulbs of infants, children, teens and adults ≤ 40 years in Metropolitan Mexico City. APOE4 carriers at higher risk of suicide accelerate their olfactory bulb pathology. Environ Res 2018; 166: 348–62.

131 Abolhasani E, Hachinski V, Ghazaleh N, Azarpazhooh MR, Mokhber N, Martin J. Air Pollution and Incidence of Dementia: A Systematic Review and Meta-analysis. Neurology 2023; 100: e242–54.

132 Yuan A, Halabicky O, Rao H, Liu J. Lifetime air pollution exposure, cognitive deficits, and brain imaging outcomes: A systematic review. NeuroToxicology 2023; 96: 69–80.

133 Zhang B, Weuve J, Langa KM, et al. Comparison of Particulate Air Pollution From Different Emission Sources and Incident Dementia in the US. JAMA Intern Med 2023; 183: 1080–9.

134 Pecheva D, Iversen JR, Palmer CE, et al. Multimodal Image Normalisation Tool (MINT) for the Adolescent Brain and Cognitive Development study: the MINT ABCD Atlas. 2022; : 2022.08.09.503395.

135 Bekierski D, Kostyrko KB, Bekierski D, Kostyrko KB. The Influence of Outdoor Particulate Matter PM2.5 on Indoor Air Quality: The Implementation of a New Assessment Method. Energies 2021; 14. DOI:10.3390/en14196230.

136 Nibagwire D, Ana GREE, Kalisa E, Twagirayezu G, Safari Kagabo A, Nsengiyumva J. Analysis of the influence of exogenous factors on indoor air quality in residential buildings. Front Built Environ 2025; 11. DOI:10.3389/fbuil.2025.1528453.

