## Supplemental Figures and Tables for "Ambient Pollution Components and Sources are Associated with Hippocampal Architecture and Memory in Pre-Adolescents"

**Supplemental Table 1.** Demographic characteristics across samples and overall ABCD sample.

| | **Overall (N=11867)** | **RSI (N=7940)** | **Long-Axis Volume (N=6795)** |
|---|---|---|---|
| **Age (months)** | | | |
| Mean (SD) | 120 (± 7.5) | 120 (± 7.4) | 120 (± 7.4) |
| Missing | <10 (<1%) | 0 (0%) | 0 (0%) |
| **Assigned Sex at Birth** | | | |
| Female | 6188 (52 %) | 4114 (52 %) | 3512 (52 %) |
| Male | 5676 (48 %) | 3826 (48 %) | 3283 (48 %) |
| Missing | <10 (<1%) | 0 (0%) | 0 (0%) |
| **Race & Ethnicity** | | | |
| non-Hispanic White | 6172 (52 %) | 4162 (52 %) | 3719 (55 %) |
| non-Hispanic Black | 1784 (15 %) | 1110 (14 %) | 882 (13 %) |
| Hispanic | 2410 (20 %) | 1676 (21 %) | 1362 (20 %) |
| Asian/Other | 1499 (13 %) | 992 (12 %) | 832 (12 %) |
| Missing | <10 (<1%) | 0 (0%) | 0 (0%) |
| **Overall Income** | | | |
| <50K | 3222 (27 %) | 2108 (27 %) | 1739 (26 %) |
| ≥50K and <100K | 3067 (26 %) | 2065 (26 %) | 1826 (27 %) |
| ≥100K | 4561 (38 %) | 3103 (39 %) | 2680 (39 %) |
| Don't know/Refuse to answer | 1015 (9 %) | 664 (8 %) | 550 (8 %) |
| Missing | <10 (<1%) | 0 (0%) | 0 (0%) |
| **Parental Highest Education** | | | |
| < HS Diploma | 593 (5 %) | 382 (5 %) | 272 (4 %) |
| HS Diploma/GED | 1132 (10 %) | 729 (9 %) | 574 (8 %) |
| Some College | 3074 (26 %) | 2041 (26 %) | 1762 (26 %) |
| Bachelor Degree | 3012 (25 %) | 1999 (25 %) | 1755 (26 %) |
| Post Graduate Degree | 4042 (34 %) | 2781 (35 %) | 2425 (36 %) |
| Missing/Refused | 14 (<1 %) | <10 (<1 %) | <10 (<1 %) |
| **MRI Manufacturer** | | | |
| GE MEDICAL SYSTEMS | 2975 (25 %) | 1870 (24 %) | 1465 (22 %) |
| Philips Medical Systems | 1523 (13 %) | 891 (11 %) | 679 (10 %) |
| SIEMENS | 7273 (61 %) | 5179 (65 %) | 4651 (68 %) |
| Missing | 96 (<1%) | 0 (0%) | 0 (0%) |

| | | | |
|---|---|---|---|
| **Proximity to Road** | | | |
| Mean (SD) | 1200 (± 1300) | 1200 (± 1200) | 1200 (± 1300) |
| Missing | 651 (5.5%) | 0 (0%) | 0 (0%) |
| **Urban vs Rural** | | | |
| Urbanized Area | 9850 (83 %) | 7036 (89 %) | 5978 (88 %) |
| Urban Clusters | 372 (3 %) | 242 (3 %) | 221 (3 %) |
| Rural | 964 (8 %) | 662 (8 %) | 596 (9 %) |
| Missing | 681 (5.7%) | 0 (0%) | 0 (0%) |
| **Neighborhood Safety** | | | |
| Mean (SD) | 3.9 (± 0.98) | 3.9 (± 0.97) | 3.9 (± 0.95) |
| Missing | <10 (0.1%) | 0 (0%) | 0 (0%) |

**Supplemental Table 2**

| Variable | Missing | % Missing |
|---|---|---|
| **Urban vs Rural** | 681 | 5.74 |
| **Proximity to Road** | 651 | 5.49 |
| **PM2.5** | 651 | 5.49 |
| **NO2** | 651 | 5.49 |
| **O3** | 651 | 5.49 |
| **Br** | 641 | 5.40 |
| **Ca** | 641 | 5.40 |
| **Cu** | 641 | 5.40 |
| **EC** | 641 | 5.40 |
| **Fe** | 641 | 5.40 |
| **K** | 641 | 5.40 |
| **$NH_4^+$** | 641 | 5.40 |
| **Ni** | 641 | 5.40 |
| **NO3** | 641 | 5.40 |
| **OC** | 641 | 5.40 |
| **PB** | 641 | 5.40 |
| **Si** | 641 | 5.40 |
| **$SO_4^{2-}$** | 641 | 5.40 |
| **V** | 641 | 5.40 |
| **Zn** | 641 | 5.40 |

| | | |
|---|---|---|
| **Race/Ethnicity** | 14 | 0.118 |
| **Neighborhood Safety** | 8 | 0.067 |
| **Sex at birth** | 3 | 0.025 |
| **Household Income** | 2 | 0.016 |
| **Age** | 1 | 0.008 |
| **Site** | 0 | 0 |
| **Highest Parental Education** | 0 | 0 |

**Supplemental Table 3.** Demographic characteristics across those included and excluded due to missing data

| | RSI Included (N=7940) | RSI Excluded (N=3927) | Long-Axis Volume Included (N=6795) | Long-Axis Volume Excluded (N=5072) |
|---|---|---|---|---|
| **Values are N (%) for categorical variables and mean (SD) for continuous variables.** | | | | |
| **Age (months)** | | | | |
| Mean (SD) | 120 (± 7.4) | 120 (± 7.7) | 120 (± 7.4) | 120 (± 7.6) |
| Missing | 0 (0%) | 1 (0.0%) | 0 (0%) | 1 (0.0%) |
| **Assigned Sex at Birth** | | | | |
| Female | 4114 (52 %) | 2074 (53 %) | 3512 (52 %) | 2676 (53 %) |
| Male | 3826 (48 %) | 1850 (47 %) | 3283 (48 %) | 2393 (47 %) |
| Intersex-Male | 0 (0 %) | 0 (0 %) | 0 (0 %) | 0 (0 %) |
| Missing | 0 (0%) | 3 (0.1%) | 0 (0%) | 3 (0.1%) |
| **Race & Ethnicity** | | | | |
| White | 4162 (52 %) | 2010 (51 %) | 3719 (55 %) | 2453 (48 %) |
| Black | 1110 (14 %) | 674 (17 %) | 882 (13 %) | 902 (18 %) |
| Hispanic | 1676 (21 %) | 734 (19 %) | 1362 (20 %) | 1048 (21 %) |
| Asian/Other | 992 (12 %) | 507 (13 %) | 832 (12 %) | 667 (13 %) |
| Missing | 0 (0%) | 2 (0.1%) | 0 (0%) | 2 (0.0%) |
| **Overall Income** | | | | |
| [<50K] | 2108 (27 %) | 1114 (28 %) | 1739 (26 %) | 1483 (29 %) |
| [≥50K and <100K] | 2065 (26 %) | 1002 (26 %) | 1826 (27 %) | 1241 (24 %) |
| [≥100K] | 3103 (39 %) | 1458 (37 %) | 2680 (39 %) | 1881 (37 %) |
| Don't know/Refuse to answer | 664 (8 %) | 351 (9 %) | 550 (8 %) | 465 (9 %) |
| Missing | 0 (0%) | <10 (0.1%) | 0 (0%) | <10 (0.0%) |
| **Parental Highest Education** | | | | |
| < HS Diploma | 382 (5 %) | 211 (5 %) | 272 (4 %) | 321 (6 %) |
| HS Diploma/GED | 729 (9 %) | 403 (10 %) | 574 (8 %) | 558 (11 %) |
| Some College | 2041 (26 %) | 1033 (26 %) | 1762 (26 %) | 1312 (26 %) |
| Bachelor Degree | 1999 (25 %) | 1013 (26 %) | 1755 (26 %) | 1257 (25 %) |
| Post Graduate Degree | 2781 (35 %) | 1261 (32 %) | 2425 (36 %) | 1617 (32 %) |
| Missing/Refused | <10 (<1 %) | <10 (<1 %) | <10 (<1 %) | <10 (<1 %) |

| | | | | |
|---|---|---|---|---|
| **MRI Manufacturer** | | | | |
| GE Medical Systems | 1870 (24 %) | 1105 (28 %) | 1465 (22 %) | 1510 (30 %) |
| Philips Medical Systems | 891 (11 %) | 632 (16 %) | 679 (10 %) | 844 (17 %) |
| SIEMENS | 5179 (65 %) | 2094 (53 %) | 4651 (68 %) | 2622 (52 %) |
| Missing | 0 (0%) | 96 (2.4%) | 0 (0%) | 96 (1.9%) |
| **Proximity to Road** | | | | |
| Mean (SD) | 1200 (± 1200) | 1200 (± 1400) | 1200 (± 1300) | 1200 (± 1300) |
| Missing | 0 (0%) | 651 (16.6%) | 0 (0%) | 651 (12.8%) |
| **Urban vs Rural** | | | | |
| Urbanized Area | 7036 (89 %) | 2814 (72 %) | 5978 (88 %) | 3872 (76 %) |
| Urban Clusters | 242 (3 %) | 130 (3 %) | 221 (3 %) | 151 (3 %) |
| Rural | 662 (8 %) | 302 (8 %) | 596 (9 %) | 368 (7 %) |
| Missing | 0 (0%) | 681 (17.3%) | 0 (0%) | 681 (13.4%) |
| **Neighborhood Safety** | | | | |
| Mean (SD) | 3.9 (± 0.97) | 3.9 (± 0.98) | 3.9 (± 0.95) | 3.9 (± 1.0) |
| Missing | 0 (0%) | 8 (0.2%) | 0 (0%) | 8 (0.2%) |

**Supplemental Table 4.** Mean and standard deviation (SD) of restriction spectrum imaging (RSI) indices and longitudinal axis volumes of the hippocampus. Abbreviations: L: Left hemisphere; R: Right hemisphere; RNT: restricted normalized total fraction, HNT: hindered normalized total fraction, FNI: right free normalized isotropic fraction.

| | **RSI (unitless) (N=7940)** |
|---|---|
| **L RNT** | |
| Mean (SD) | 0.36 (± 0.022) |
| **L HNT** | |
| Mean (SD) | 0.78 (± 0.031) |
| **L FNI** | |
| Mean (SD) | 0.37 (± 0.044) |
| **R RNT** | |
| Mean (SD) | 0.36 (± 0.021) |
| **R HNT** | |
| Mean (SD) | 0.78 (± 0.030) |
| **R FNI** | |
| Mean (SD) | 0.37 (± 0.041) |
| | **Long-Axis Volume ($mm^3$) (N=6795)** |
| **L Head** | |
| Mean (SD) | 1700 (± 210) |
| **L Body** | |
| Mean (SD) | 1100 (± 130) |
| **L Tail** | |
| Mean (SD) | 530 (± 71) |
| **R Head** | |
| Mean (SD) | 1700 (± 210) |
| **R Body** | |
| Mean (SD) | 1100 (± 120) |
| **R Tail** | |
| Mean (SD) | 530 (± 73) |

**Supplemental Table 5.** Descriptive statistics of the air pollution metrics. We assigned 2016 annual average air pollution exposure, $PM_{2.5}$ (μg/m³), $NO_2$ (ppb), and 8-hour maximum ground-level $O_3$ (ppb), to each child's primary residence collected from the caregiver during the initial ABCD Study enrollment visits (2016-2018). To estimate residential exposure to $PM_{2.5}$ constituent components, similar machine learning-based models were used to calculate 15 components based on component measurements available. EC, $NH_4^+$, $NO_3$, OC, and $SO_4^{2-}$ were all originally in μg/m³ and were converted to ng/m³ prior to analysis to ensure all pollutants were on the same scale. Abbreviations: Br: bromine, Ca: calcium, Cu: copper, EC: elemental carbon, Fe: iron, K: potassium, $NH_4^+$: ammonium, Ni: nickel, $NO_2$: nitrogen dioxide, $NO_3$: nitrate, $O_3$: ozone, OC: organic carbon, Pb: lead, $PM_{2.5}$: fine particulate matter, Si: silicon, $SO_4^{2-}$: sulfate, V: vanadium, Zn: zinc

| | **Overall (N=11867)** | **RSI (N=7940)** | **Long-Axis Volume (N=6795)** |
|---|---|---|---|
| **PM2.5 (μg/m3)** | | | |
| Mean (SD) | 7.7 (± 1.6) | 7.7 (± 1.6) | 7.6 (± 1.6) |
| Missing | 651 (5.5%) | 0 (0%) | 0 (0%) |
| **NO2 (ppb)** | | | |
| Mean (SD) | 19 (± 5.8) | 19 (± 5.7) | 18 (± 5.6) |
| Missing | 651 (5.5%) | 0 (0%) | 0 (0%) |
| **O3 (8-hr max, ppb)** | | | |
| Mean (SD) | 42 (± 4.4) | 42 (± 4.4) | 42 (± 4.4) |
| Missing | 651 (5.5%) | 0 (0%) | 0 (0%) |
| | *ng/m³* | *ng/m³* | *ng/m³* |
| **Br** | | | |
| Mean (SD) | 2.6 (± 0.58) | 2.6 (± 0.60) | 2.6 (± 0.58) |
| Missing | 641 (5.4%) | 0 (0%) | 0 (0%) |
| **Ca** | | | |
| Mean (SD) | 48 (± 21) | 48 (± 21) | 48 (± 22) |
| Missing | 641 (5.4%) | 0 (0%) | 0 (0%) |
| **Cu** | | | |
| Mean (SD) | 4.6 (± 1.7) | 4.6 (± 1.7) | 4.5 (± 1.7) |
| Missing | 641 (5.4%) | 0 (0%) | 0 (0%) |
| **Fe** | | | |
| Mean (SD) | 66 (± 25) | 66 (± 25) | 65 (± 24) |
| Missing | 641 (5.4%) | 0 (0%) | 0 (0%) |
| **K** | | | |
| Mean (SD) | 63 (± 9.8) | 63 (± 9.9) | 63 (± 9.7) |
| Missing | 641 (5.4%) | 0 (0%) | 0 (0%) |
| **Ni** | | | |
| Mean (SD) | 0.83 (± 0.27) | 0.84 (± | 0.82 (± 0.27) |

| | | | |
|---|---|---|---|
| | | 0.27) | |
| Missing | 641 (5.4%) | 0 (0%) | 0 (0%) |
| **Pb** | | | |
| Mean (SD) | 4.5 (± 1.2) | 4.5 (± 1.2) | 4.5 (± 1.3) |
| Missing | 641 (5.4%) | 0 (0%) | 0 (0%) |
| **Si** | | | |
| Mean (SD) | 84 (± 38) | 84 (± 38) | 85 (± 38) |
| Missing | 641 (5.4%) | 0 (0%) | 0 (0%) |
| **V** | | | |
| Mean (SD) | 0.37 (± 0.20) | 0.37 (± 0.20) | 0.38 (± 0.21) |
| Missing | 641 (5.4%) | 0 (0%) | 0 (0%) |
| **Zn** | | | |
| Mean (SD) | 9.3 (± 4.1) | 9.3 (± 4.1) | 9.1 (± 4.1) |
| Missing | 641 (5.4%) | 0 (0%) | 0 (0%) |
| **EC** | | | |
| Mean (SD) | 0.52 (± 0.16) | 530 (± 160) | 520 (± 150) |
| Missing | 641 (5.4%) | 0 (0%) | 0 (0%) |
| **$NH_4^+$** | | | |
| Mean (SD) | 0.30 (± 0.12) | 300 (± 120) | 280 (± 120) |
| Missing | 641 (5.4%) | 0 (0%) | 0 (0%) |
| **$NO_3$** | | | |
| Mean (SD) | 0.93 (± 0.35) | 920 (± 360) | 900 (± 350) |
| Missing | 641 (5.4%) | 0 (0%) | 0 (0%) |
| **OC** | | | |
| Mean (SD) | 1.9 (± 0.45) | 1900 (± 460) | 1900 (± 460) |
| Missing | 641 (5.4%) | 0 (0%) | 0 (0%) |
| **$SO_4^{2-}$** | | | |
| Mean (SD) | 0.92 (± 0.29) | 920 (± 290) | 890 (± 290) |
| Missing | 641 (5.4%) | 0 (0%) | 0 (0%) |

**Supplemental Table 6.** Learning and memory was measured using the Rey Auditory Verbal Learning Test. The RAVLT evaluates verbal memory by presenting a list of 15 unrelated words (List A) over five learning trials, with participants recalling as many words as possible after each trial. Following these trials, a second list of 15 words (List B) is introduced as a proactive mnemonic interference task, after which participants immediately recall List A. Delayed recall of List A was then assessed 20–30 minutes later. Our primary outcomes included: learning (the difference in words recalled between Trial 5 and Trial 1), proactive mnemonic interference (total words recalled from List B), immediate recall (total words recalled from List A immediately after interference), and delayed recall (total words recalled from List A after the delay)

| | **Overall (N=11493)** | **RAVLT (N=7699)** |
|---|---|---|
| **Learning (Trial V - Trial I)** | | |
| Mean (SD) | 6.2 (± 2.5) | 6.2 (± 2.5) |
| **Immediate Recall** | | |
| Mean (SD) | 9.7 (± 3.0) | 9.8 (± 3.0) |
| **Proactive Interference (List B)** | | |
| Mean (SD) | 4.9 (± 1.7) | 4.9 (± 1.7) |
| **Delayed Recall** | | |
| Mean (SD) | 9.2 (± 3.2) | 9.3 (± 3.2) |

**Supplemental Table 7.** Only latent dimensions from significant PLSC models are interpreted. The number of extractable latent factors is limited by the smaller variable set between blocks (hippocampal RSI vs. PM2.5 total mass/sources/components). Significant latent dimensions are bolded.

| **Partial Least Squares Correlation Model Statistics and Eigenvalues for Hippocampal RSI and Air Pollution Models** | | | | | |
|---|---|---|---|---|---|
| **PM2.5 Total Mass** | | **PM2.5 Components** | | **PM2.5 Sources** | |
| *p(model)* | .0001 | .0001 | | .0001 | |
| Latent Dimension | Proportion of Variance, τ | Latent Dimension | Proportion of Variance, τ | Latent Dimension | Proportion of Variance, τ |
| 1 | **81%** | 1 | **72%** | 1 | **61%** |
| 2 | 17% | 2 | **24%** | 2 | **32%** |
| 3 | 2% | 3 | 2% | 3 | 5% |
| - | - | 4 | 1% | 4 | 1% |
| - | - | 5 | 1% | 5 | 1% |
| - | - | 6 | 0% | 6 | 0% |

**Supplemental Table 8.** Only latent dimensions from significant PLSC models are interpreted. The number of extractable latent factors is limited by the smaller variable set between blocks (hippocampal long-axis volumes vs. PM2.5 total mass/sources/components).

**Partial Least Squares Correlation Model Statistics and Eigenvalues for Hippocampal Long-Axis and Air Pollution Models**

| **PM2.5 Total Mass** | | **PM2.5 Components** | | **PM2.5 Sources** | |
|---|---|---|---|---|---|
| *p(model)* | .98 | .0004 | | .041 | |
| Latent Dimension | Proportion of Variance, *τ* | Latent Dimension | Proportion of Variance, *τ* | Latent Dimension | Proportion of Variance, *τ* |
| 1 | 67% | 1 | **75%** | 1 | **77%** |
| 2 | 24% | 2 | 14% | 2 | 12% |
| 3 | 9% | 3 | 5% | 3 | 5% |
| - | - | 4 | 4% | 4 | 3% |
| - | - | 5 | 2% | 5 | 3% |
| - | - | 6 | 1% | 6 | 0% |

**Supplemental Table 9.** Only latent dimensions from significant PLSC models are interpreted. The number of extractable latent factors is limited by the smaller variable set between blocks (RAVLT performance vs. PM2.5 total mass/sources/components). Significant latent dimensions are bolded.

| **Partial Least Squares Correlation Model Statistics and Eigenvalues for RAVLT Performance and Air Pollution Models** | | | | | |
|---|---|---|---|---|---|
| **PM2.5 Total Mass** | | **PM2.5 Components** | | **PM2.5 Sources** | |
| *p(model)* | .96 | .0001 | | .002 | |
| Latent Dimension | Proportion of Variance, *τ* | Latent Dimension | Proportion of Variance, *τ* | Latent Dimension | Proportion of Variance, *τ* |
| 1 | 66% | 1 | **66.5%** | 1 | **58.24%** |
| 2 | 31% | 2 | **24.1%** | 2 | **38.18%** |
| 3 | 3% | 3 | 8.4% | 3 | 3.2% |
| - | - | 4 | 1% | 4 | .38% |

**Supplemental Table 10.** Only latent dimensions from significant PLSC models are interpreted. The number of extractable latent factors is limited by the smaller variable set between blocks (hippocampal architecture vs. RAVLT performance). Significant latent dimensions are bolded.

| **Partial Least Squares Correlation Model Statistics and Eigenvalues for RAVLT Performance and Hippocampal Architecture** | | | |
|---|---|---|---|
| **Hippocampal RSI (RNT, HNT, FNI)** | | **Hippocampus Long-Axis** | |
| *p(model)* | .0001 | .0001 | |
| Latent Dimension | Proportion of Variance, $\tau$ | Latent Dimension | Proportion of Variance, $\tau$ |
| 1 | **82%** | 1 | **96%** |
| 2 | 16**%** | 2 | 3% |
| 3 | 1% | 3 | 1% |
| 4 | 0% | 4 | 0% |

**Supplemental Figure 1.** Flowchart for participant selection for hippocampal restriction spectrum imaging (RSI) and long-axis volume analyses.

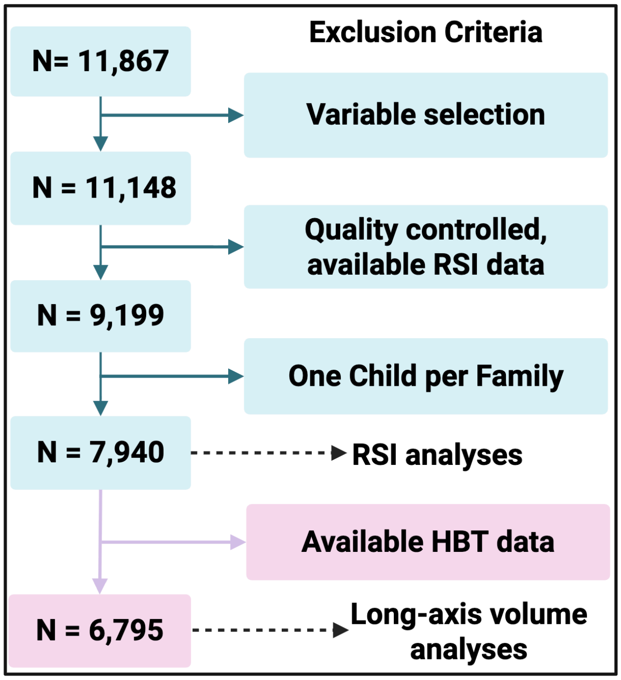

**Supplemental Figure 2.** Spearman correlations of exposures to criteria pollutants ($PM_{2.5}$ total mass, $NO_2$, and $O_3$), fifteen $PM_{2.5}$components, and six sources of $PM_{2.5}$ for the RSI sample (N=7,940). Abbreviations: Br: bromine, Ca: calcium, Cu: copper, EC: elemental carbon, Fe: iron, K: potassium, NH4: ammonium, Ni: nickel, $NO_2$: nitrogen dioxide, $NO_3$: nitrate, $O_3$: ozone, OC: organic carbon, Pb: lead, $PM_{2.5}$: fine particulate matter, Si: silicon, SO4: sulfate, V: vanadium, Zn: zinc

**Spearman Correlation**

| | Br | Cu | Fe | EC | OC | NH4 | NO3 | SO4 | V | Zn | Ni | Pb | Ca | Si | K | PM2.5 | O3 | NO2 | Crustal | Sulfates | Biomass | Traffic | Nitrates | Industrial |
|---|---|---|---|---|---|---|---|---|---|---|---|---|---|---|---|---|---|---|---|---|---|---|---|---|
| Br | 1.00 | 0.51 | 0.63 | 0.49 | 0.40 | 0.31 | 0.47 | 0.35 | 0.32 | 0.30 | 0.33 | 0.12 | 0.59 | 0.55 | 0.59 | 0.57 | -0.05 | 0.16 | 0.43 | 0.13 | 0.34 | 0.44 | 0.43 | -0.07 |
| Cu | 0.51 | 1.00 | 0.83 | 0.77 | 0.50 | 0.24 | 0.41 | 0.26 | 0.25 | 0.53 | 0.58 | 0.39 | 0.40 | 0.25 | 0.45 | 0.52 | -0.01 | 0.43 | 0.21 | -0.01 | 0.06 | 0.91 | 0.30 | 0.27 |
| Fe | 0.63 | 0.83 | 1.00 | 0.82 | 0.49 | 0.25 | 0.51 | 0.25 | 0.30 | 0.50 | 0.50 | 0.27 | 0.68 | 0.58 | 0.55 | 0.59 | -0.08 | 0.48 | 0.48 | -0.09 | 0.20 | 0.84 | 0.40 | 0.07 |
| EC | 0.49 | 0.77 | 0.82 | 1.00 | 0.65 | 0.18 | 0.35 | 0.24 | 0.19 | 0.53 | 0.55 | 0.32 | 0.40 | 0.34 | 0.50 | 0.52 | 0.00 | 0.53 | 0.25 | -0.02 | 0.28 | 0.91 | 0.22 | 0.21 |
| OC | 0.40 | 0.50 | 0.49 | 0.65 | 1.00 | 0.19 | 0.20 | 0.27 | 0.10 | 0.40 | 0.49 | 0.15 | 0.12 | 0.19 | 0.57 | 0.40 | -0.14 | 0.20 | 0.05 | 0.12 | 0.61 | 0.63 | 0.03 | 0.14 |
| NH4 | 0.31 | 0.24 | 0.25 | 0.18 | 0.19 | 1.00 | 0.62 | 0.57 | -0.15 | 0.50 | 0.29 | 0.40 | 0.18 | -0.05 | -0.03 | 0.45 | -0.30 | 0.19 | -0.25 | 0.37 | -0.23 | 0.17 | 0.74 | 0.31 |
| NO3 | 0.47 | 0.41 | 0.51 | 0.35 | 0.20 | 0.62 | 1.00 | 0.01 | -0.16 | 0.57 | 0.30 | 0.36 | 0.37 | 0.18 | 0.10 | 0.32 | -0.21 | 0.45 | -0.10 | -0.32 | 0.21 | 0.31 | 0.95 | 0.25 |
| SO4 | 0.35 | 0.26 | 0.25 | 0.24 | 0.27 | 0.57 | 0.01 | 1.00 | 0.30 | 0.08 | 0.26 | 0.04 | 0.24 | 0.13 | 0.30 | 0.58 | -0.13 | -0.12 | 0.13 | 0.89 | -0.30 | 0.28 | 0.09 | -0.05 |
| V | 0.32 | 0.25 | 0.30 | 0.19 | 0.10 | -0.15 | -0.16 | 0.30 | 1.00 | -0.03 | 0.25 | 0.02 | 0.45 | 0.44 | 0.43 | 0.31 | -0.05 | -0.05 | 0.55 | 0.27 | -0.10 | 0.22 | -0.20 | -0.06 |
| Zn | 0.30 | 0.53 | 0.50 | 0.53 | 0.40 | 0.50 | 0.57 | 0.08 | -0.03 | 1.00 | 0.54 | 0.76 | 0.14 | -0.08 | 0.13 | 0.34 | -0.16 | 0.55 | -0.24 | -0.13 | -0.02 | 0.53 | 0.60 | 0.76 |
| Ni | 0.33 | 0.58 | 0.50 | 0.55 | 0.49 | 0.29 | 0.30 | 0.26 | 0.25 | 0.54 | 1.00 | 0.56 | 0.09 | 0.01 | 0.17 | 0.36 | -0.07 | 0.35 | -0.06 | 0.14 | 0.01 | 0.62 | 0.21 | 0.54 |
| Pb | 0.12 | 0.39 | 0.27 | 0.32 | 0.15 | 0.40 | 0.36 | 0.04 | 0.02 | 0.76 | 0.56 | 1.00 | 0.01 | -0.20 | -0.11 | 0.20 | -0.17 | 0.48 | -0.26 | -0.04 | -0.31 | 0.31 | 0.41 | 0.93 |
| Ca | 0.59 | 0.40 | 0.68 | 0.40 | 0.12 | 0.18 | 0.37 | 0.24 | 0.45 | 0.14 | 0.09 | 0.01 | 1.00 | 0.84 | 0.54 | 0.52 | -0.20 | 0.22 | 0.84 | -0.04 | 0.07 | 0.29 | 0.34 | -0.24 |
| Si | 0.55 | 0.25 | 0.58 | 0.34 | 0.19 | -0.05 | 0.18 | 0.13 | 0.44 | -0.08 | 0.01 | -0.20 | 0.84 | 1.00 | 0.59 | 0.38 | -0.12 | 0.11 | 0.89 | -0.06 | 0.33 | 0.22 | 0.09 | -0.44 |
| K | 0.59 | 0.45 | 0.55 | 0.50 | 0.57 | -0.03 | 0.10 | 0.30 | 0.43 | 0.13 | 0.17 | -0.11 | 0.54 | 0.59 | 1.00 | 0.48 | -0.09 | 0.00 | 0.57 | 0.13 | 0.49 | 0.43 | -0.05 | -0.21 |
| PM2.5 | 0.57 | 0.52 | 0.59 | 0.52 | 0.40 | 0.45 | 0.32 | 0.58 | 0.31 | 0.34 | 0.36 | 0.20 | 0.52 | 0.38 | 0.48 | 1.00 | -0.21 | 0.22 | 0.32 | 0.36 | -0.01 | 0.49 | 0.32 | 0.05 |
| O3 | -0.05 | -0.01 | -0.08 | 0.00 | -0.14 | -0.30 | -0.21 | -0.13 | -0.05 | -0.16 | -0.07 | -0.17 | -0.20 | -0.12 | -0.09 | -0.21 | 1.00 | -0.02 | -0.06 | -0.04 | 0.01 | 0.05 | -0.22 | -0.12 |
| NO2 | 0.16 | 0.43 | 0.48 | 0.53 | 0.20 | 0.19 | 0.45 | -0.12 | -0.05 | 0.55 | 0.35 | 0.48 | 0.22 | 0.11 | 0.00 | 0.22 | -0.02 | 1.00 | 0.00 | -0.30 | 0.04 | 0.48 | 0.39 | 0.41 |
| Crustal | 0.43 | 0.21 | 0.48 | 0.25 | 0.05 | -0.25 | -0.10 | 0.13 | 0.55 | -0.24 | -0.06 | -0.26 | 0.84 | 0.89 | 0.57 | 0.32 | -0.06 | 0.00 | 1.00 | 0.01 | 0.16 | 0.15 | -0.16 | -0.49 |
| Sulfates | 0.13 | -0.01 | -0.09 | -0.02 | 0.12 | 0.37 | -0.32 | 0.89 | 0.27 | -0.13 | 0.14 | -0.04 | -0.04 | -0.06 | 0.13 | 0.36 | -0.04 | -0.30 | 0.01 | 1.00 | -0.39 | 0.03 | -0.21 | -0.06 |
| Biomass | 0.34 | 0.06 | 0.20 | 0.28 | 0.61 | -0.23 | 0.21 | -0.30 | -0.10 | -0.02 | 0.01 | -0.31 | 0.07 | 0.33 | 0.49 | -0.01 | 0.01 | 0.04 | 0.16 | -0.39 | 1.00 | 0.16 | -0.01 | -0.32 |
| Traffic | 0.44 | 0.91 | 0.84 | 0.91 | 0.63 | 0.17 | 0.31 | 0.28 | 0.22 | 0.53 | 0.62 | 0.31 | 0.29 | 0.22 | 0.43 | 0.49 | 0.05 | 0.48 | 0.15 | 0.03 | 0.16 | 1.00 | 0.17 | 0.24 |
| Nitrates | 0.43 | 0.30 | 0.40 | 0.22 | 0.03 | 0.74 | 0.95 | 0.09 | -0.20 | 0.60 | 0.21 | 0.41 | 0.34 | 0.09 | -0.05 | 0.32 | -0.22 | 0.39 | -0.16 | -0.21 | -0.01 | 0.17 | 1.00 | 0.30 |
| Industrial | -0.07 | 0.27 | 0.07 | 0.21 | 0.14 | 0.31 | 0.25 | -0.05 | -0.06 | 0.76 | 0.54 | 0.93 | -0.24 | -0.44 | -0.21 | 0.05 | -0.12 | 0.41 | -0.49 | -0.06 | -0.32 | 0.24 | 0.30 | 1.00 |

**Supplemental Figure 3.** Spearman correlations of restriction spectrum imaging analyses participant exposures to criteria pollutants ($PM_{2.5}$ total mass, $NO_2$, and $O_3$), fifteen $PM_{2.5}$ components, and six sources of $PM_{2.5}$, $NO_2$, and $O_3$ for the long-axis sample (N=6,795). Abbreviations: Br: bromine, Ca: calcium, Cu: copper, EC: elemental carbon, Fe: iron, K: potassium, NH4: ammonium, Ni: nickel, $NO_2$: nitrogen dioxide, $NO_3$: nitrate, $O_3$: ozone, OC: organic carbon, Pb: lead, $PM_{2.5}$: fine particulate matter, Si: silicon, SO4: sulfate, V: vanadium, Zn: zinc

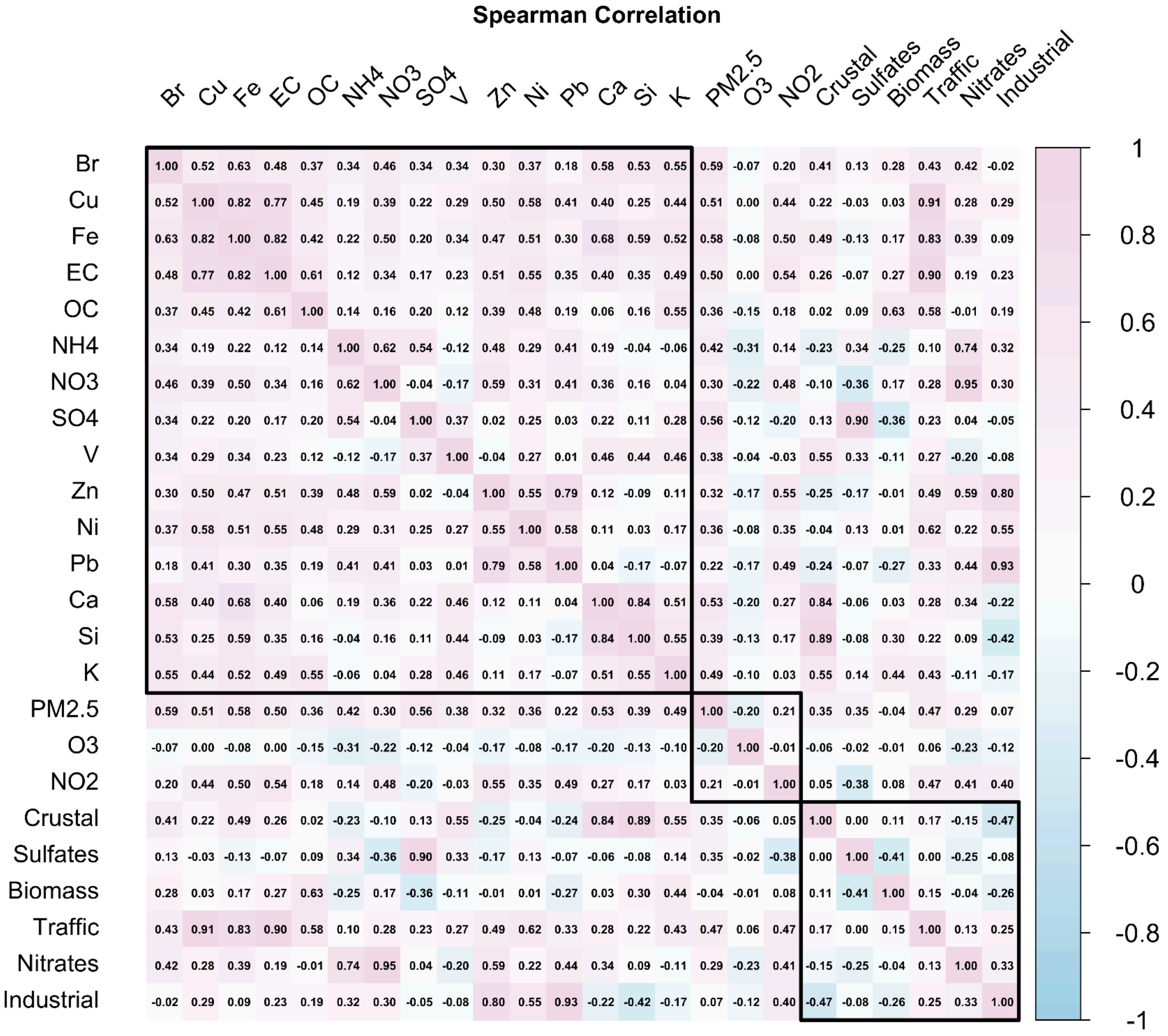
Spearman Correlation
Br Cu Fe EC OC NH4 NO3 SO4 V Zn Ni Pb Ca Si K PM2.5 O3 NO2 Crustal Sulfates Biomass Traffic Nitrates Industrial
Br 1.00 0.52 0.63 0.48 0.37 0.34 0.46 0.34 0.34 0.30 0.37 0.18 0.58 0.53 0.55 0.59 -0.07 0.20 0.41 0.13 0.28 0.43 0.42 -0.02
Cu 0.52 1.00 0.82 0.77 0.45 0.19 0.39 0.22 0.29 0.50 0.58 0.41 0.40 0.25 0.44 0.51 0.00 0.44 0.22 -0.03 0.03 0.91 0.28 0.29
Fe 0.63 0.82 1.00 0.82 0.42 0.22 0.50 0.20 0.34 0.47 0.51 0.30 0.68 0.59 0.52 0.58 -0.08 0.50 0.49 -0.13 0.17 0.83 0.39 0.09
EC 0.48 0.77 0.82 1.00 0.61 0.12 0.34 0.17 0.23 0.51 0.55 0.35 0.40 0.35 0.49 0.50 0.00 0.54 0.26 -0.07 0.27 0.90 0.19 0.23
OC 0.37 0.45 0.42 0.61 1.00 0.14 0.16 0.20 0.12 0.39 0.48 0.19 0.06 0.16 0.55 0.36 -0.15 0.18 0.02 0.09 0.63 0.58 -0.01 0.19
NH4 0.34 0.19 0.22 0.12 0.14 1.00 0.62 0.54 -0.12 0.48 0.29 0.41 0.19 -0.04 -0.06 0.42 -0.31 0.14 -0.23 0.34 -0.25 0.10 0.74 0.32
NO3 0.46 0.39 0.50 0.34 0.16 0.62 1.00 -0.04 -0.17 0.59 0.31 0.41 0.36 0.16 0.04 0.30 -0.22 0.48 -0.10 -0.36 0.17 0.28 0.95 0.30
SO4 0.34 0.22 0.20 0.17 0.20 0.54 -0.04 1.00 0.37 0.02 0.25 0.03 0.22 0.11 0.28 0.56 -0.12 -0.20 0.13 0.90 -0.36 0.23 0.04 -0.05
V 0.34 0.29 0.34 0.23 0.12 -0.12 -0.17 0.37 1.00 -0.04 0.27 0.01 0.46 0.44 0.46 0.38 -0.04 -0.03 0.55 0.33 -0.11 0.27 -0.20 -0.08
Zn 0.30 0.50 0.47 0.51 0.39 0.48 0.59 0.02 -0.04 1.00 0.55 0.79 0.12 -0.09 0.11 0.32 -0.17 0.55 -0.25 -0.17 -0.01 0.49 0.59 0.80
Ni 0.37 0.58 0.51 0.55 0.48 0.29 0.31 0.25 0.27 0.55 1.00 0.58 0.11 0.03 0.17 0.36 -0.08 0.35 -0.04 0.13 0.01 0.62 0.22 0.55
Pb 0.18 0.41 0.30 0.35 0.19 0.41 0.41 0.03 0.01 0.79 0.58 1.00 0.04 -0.17 -0.07 0.22 -0.17 0.49 -0.24 -0.07 -0.27 0.33 0.44 0.93
Ca 0.58 0.40 0.68 0.40 0.06 0.19 0.36 0.22 0.46 0.12 0.11 0.04 1.00 0.84 0.51 0.53 -0.20 0.27 0.84 -0.06 0.03 0.28 0.34 -0.22
Si 0.53 0.25 0.59 0.35 0.16 -0.04 0.16 0.11 0.44 -0.09 0.03 -0.17 0.84 1.00 0.55 0.39 -0.13 0.17 0.89 -0.08 0.30 0.22 0.09 -0.42
K 0.55 0.44 0.52 0.49 0.55 -0.06 0.04 0.28 0.46 0.11 0.17 -0.07 0.51 0.55 1.00 0.49 -0.10 0.03 0.55 0.14 0.44 0.43 -0.11 -0.17
PM2.5 0.59 0.51 0.58 0.50 0.36 0.42 0.30 0.56 0.38 0.32 0.36 0.22 0.53 0.39 0.49 1.00 -0.20 0.21 0.35 0.35 -0.04 0.47 0.29 0.07
O3 -0.07 0.00 -0.08 0.00 -0.15 -0.31 -0.22 -0.12 -0.04 -0.17 -0.08 -0.17 -0.20 -0.13 -0.10 -0.20 1.00 -0.01 -0.06 -0.02 -0.01 0.06 -0.23 -0.12
NO2 0.20 0.44 0.50 0.54 0.18 0.14 0.48 -0.20 -0.03 0.55 0.35 0.49 0.27 0.17 0.03 0.21 -0.01 1.00 0.05 -0.38 0.08 0.47 0.41 0.40
Crustal 0.41 0.22 0.49 0.26 0.02 -0.23 -0.10 0.13 0.55 -0.25 -0.04 -0.24 0.84 0.89 0.55 0.35 -0.06 0.05 1.00 0.00 0.11 0.17 -0.15 -0.47
Sulfates 0.13 -0.03 -0.13 -0.07 0.09 0.34 -0.36 0.90 0.33 -0.17 0.13 -0.07 -0.06 -0.08 0.14 0.35 -0.02 -0.38 0.00 1.00 -0.41 0.00 -0.25 -0.08
Biomass 0.28 0.03 0.17 0.27 0.63 -0.25 0.17 -0.36 -0.11 -0.01 0.01 -0.27 0.03 0.30 0.44 -0.04 -0.01 0.08 0.11 -0.41 1.00 0.15 -0.04 -0.26
Traffic 0.43 0.91 0.83 0.90 0.58 0.10 0.28 0.23 0.27 0.49 0.62 0.33 0.28 0.22 0.43 0.47 0.06 0.47 0.17 0.00 0.15 1.00 0.13 0.25
Nitrates 0.42 0.28 0.39 0.19 -0.01 0.74 0.95 0.04 -0.20 0.59 0.22 0.44 0.34 0.09 -0.11 0.29 -0.23 0.41 -0.15 -0.25 -0.04 0.13 1.00 0.33
Industrial -0.02 0.29 0.09 0.23 0.19 0.32 0.30 -0.05 -0.08 0.80 0.55 0.93 -0.22 -0.42 -0.17 0.07 -0.12 0.40 -0.47 -0.08 -0.26 0.25 0.33 1.00
1
0.8
0.6
0.4
0.2
0
-0.2
-0.4
-0.6
-0.8
-1

**Supplemental Figure 4.** Spearman correlations of restriction spectrum imaging analyses for participants (N=7,940). Abbreviations: L: Left hemisphere; R: Right hemisphere; RNT: restricted normalized total fraction, HNT: hindered normalized total fraction, FNI: right free normalized isotropic fraction.

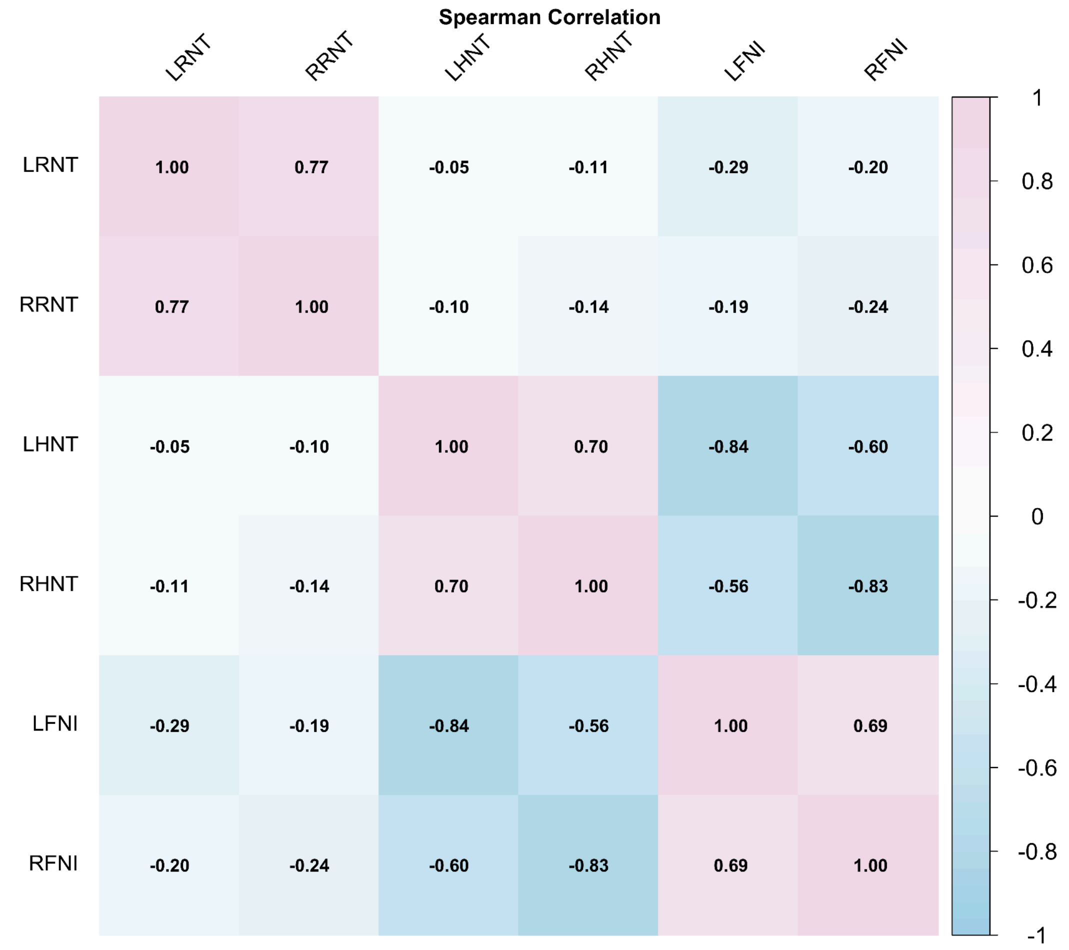

**Supplemental Figure 5.** Spearman correlations of hippocampal long-axis volumes for participants (N=6,795).

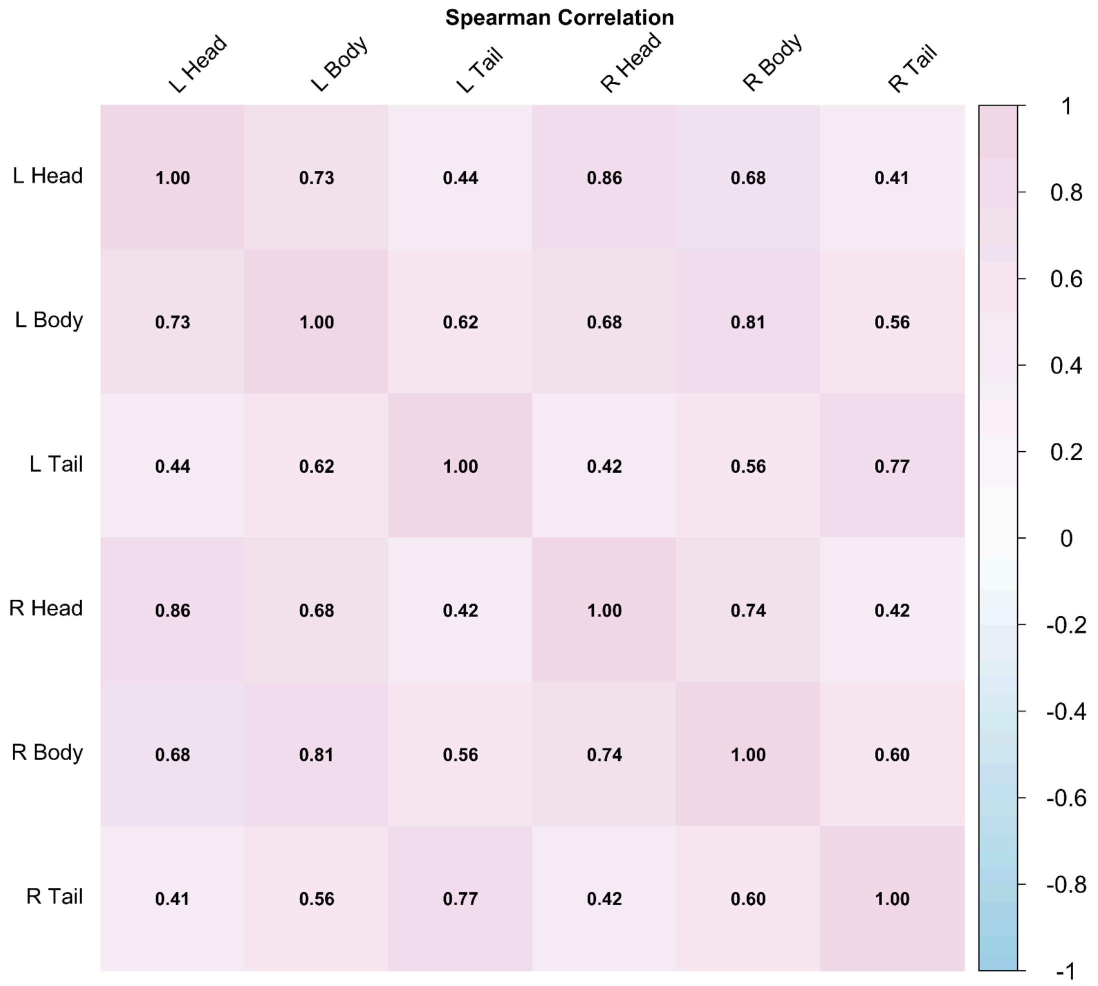

**Supplemental Figure 6.** Scree plot depicting the eigenvalues for RSI and PM2.5 total mass, $NO_2$, and $O_3$ analyses. The first latent dimension is identified as significant, indicating the primary sources of shared variance between the datasets. The elbow point suggests the optimal number of dimensions to retain in the model.

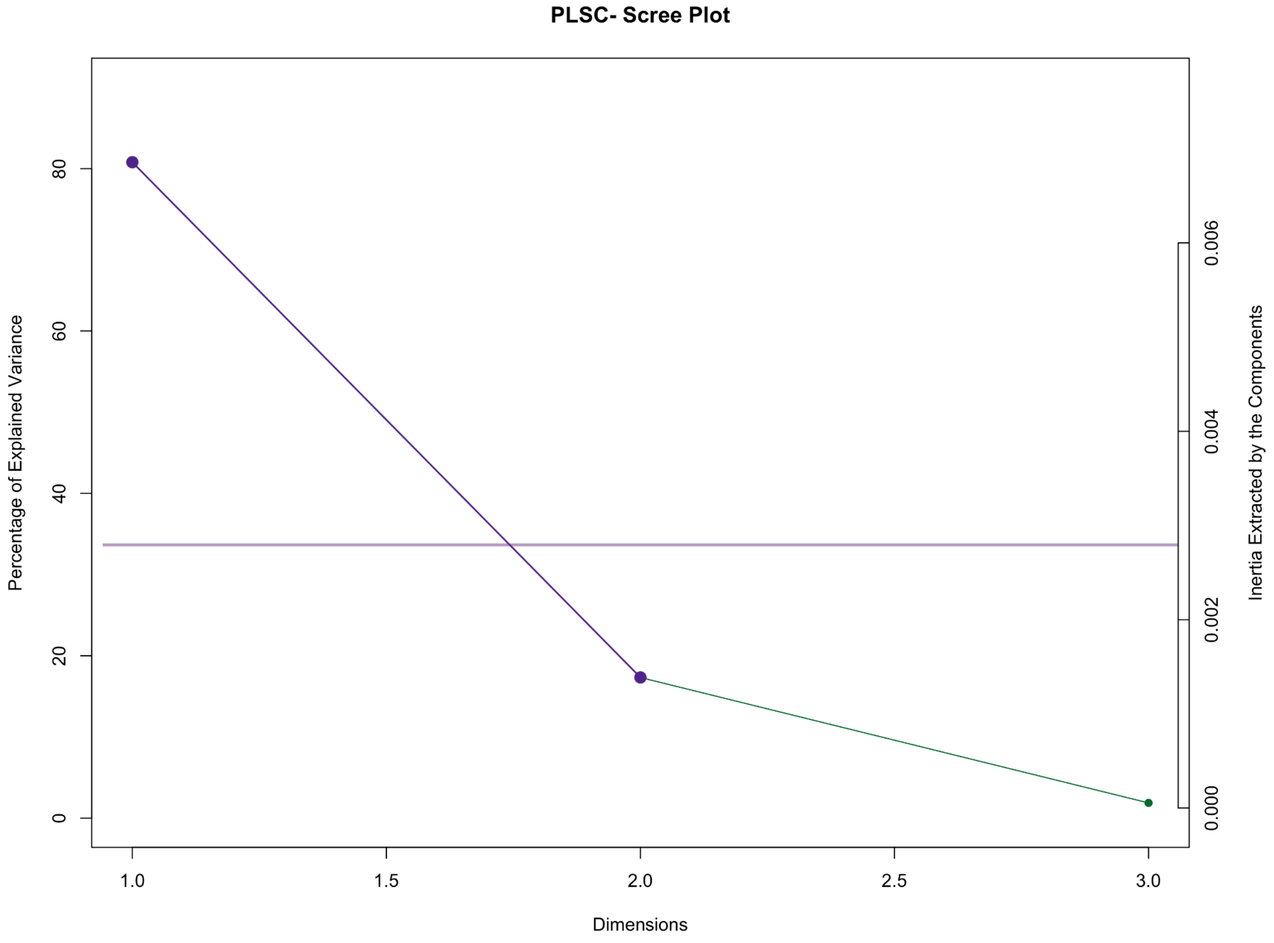

**Supplemental Figure 7.** Scree plot depicting the eigenvalues for RSI and PM2.5 component analyses. The first and second latent dimensions are identified as significant, indicating the primary sources of shared variance between the datasets. The elbow point suggests the optimal number of dimensions to retain in the model

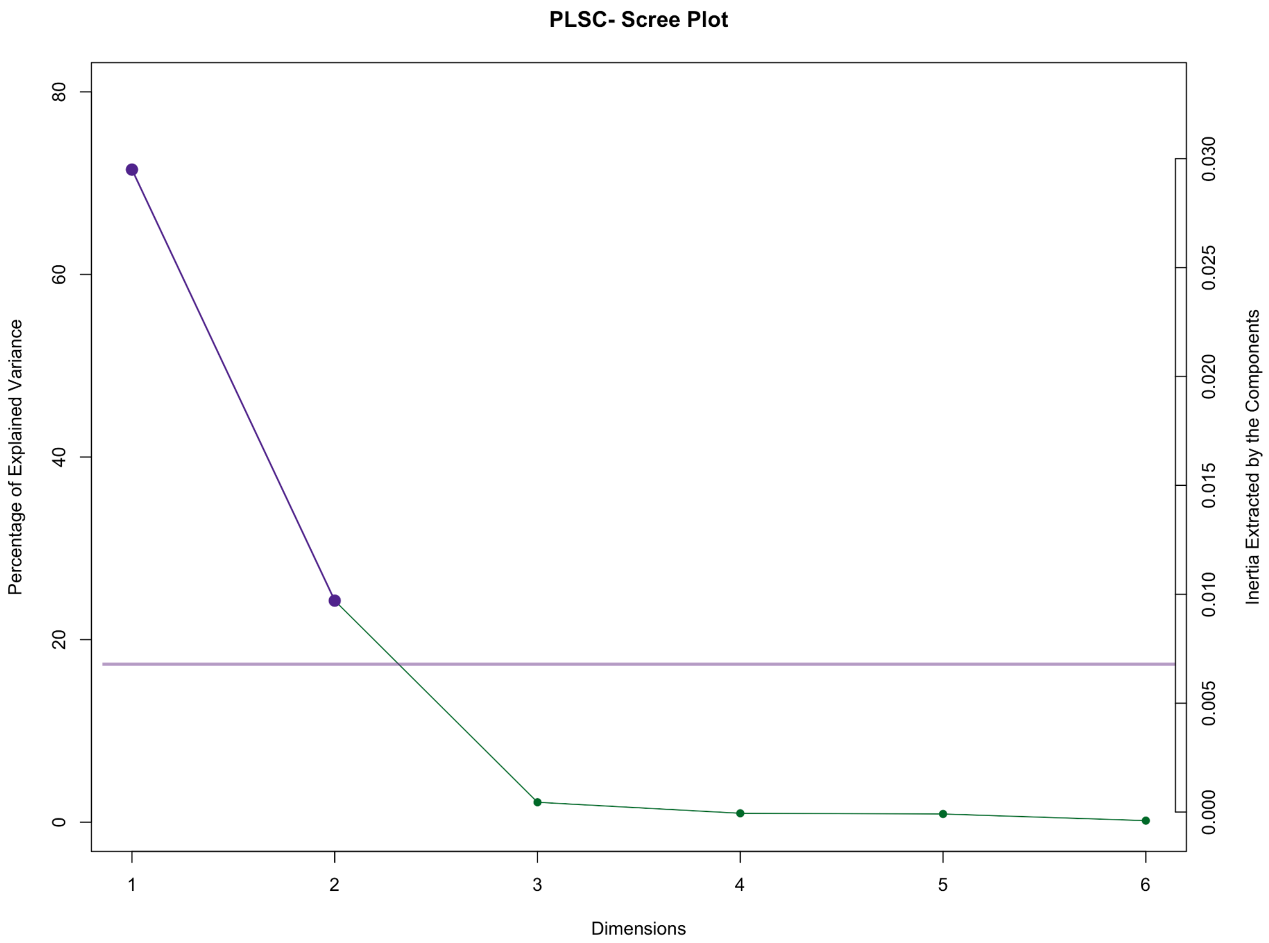

**Supplemental Figure 8.** Scree plot depicting the eigenvalues for RSI and PM2.5 source analyses. The first and second latent dimensions are identified as significant, indicating the primary sources of shared variance between the datasets. The elbow point suggests the optimal number of dimensions to retain in the model.

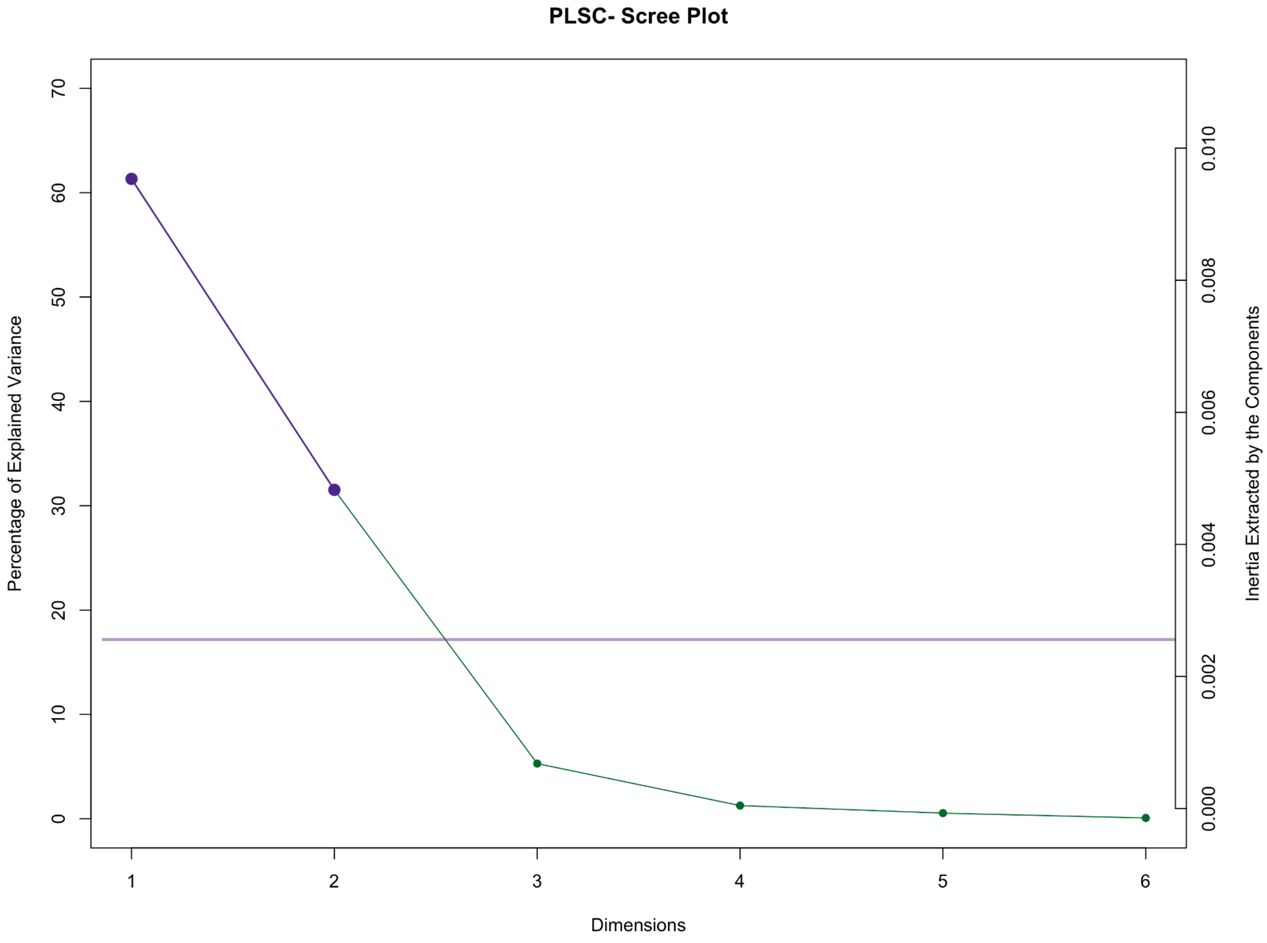

**Supplemental Figure 9.** Scree plot depicting the eigenvalues for hippocampal head, body, and tail volume and PM2.5 total mass, $NO_2$, and $O_3$ analyses. There were no significant latent dimensions.

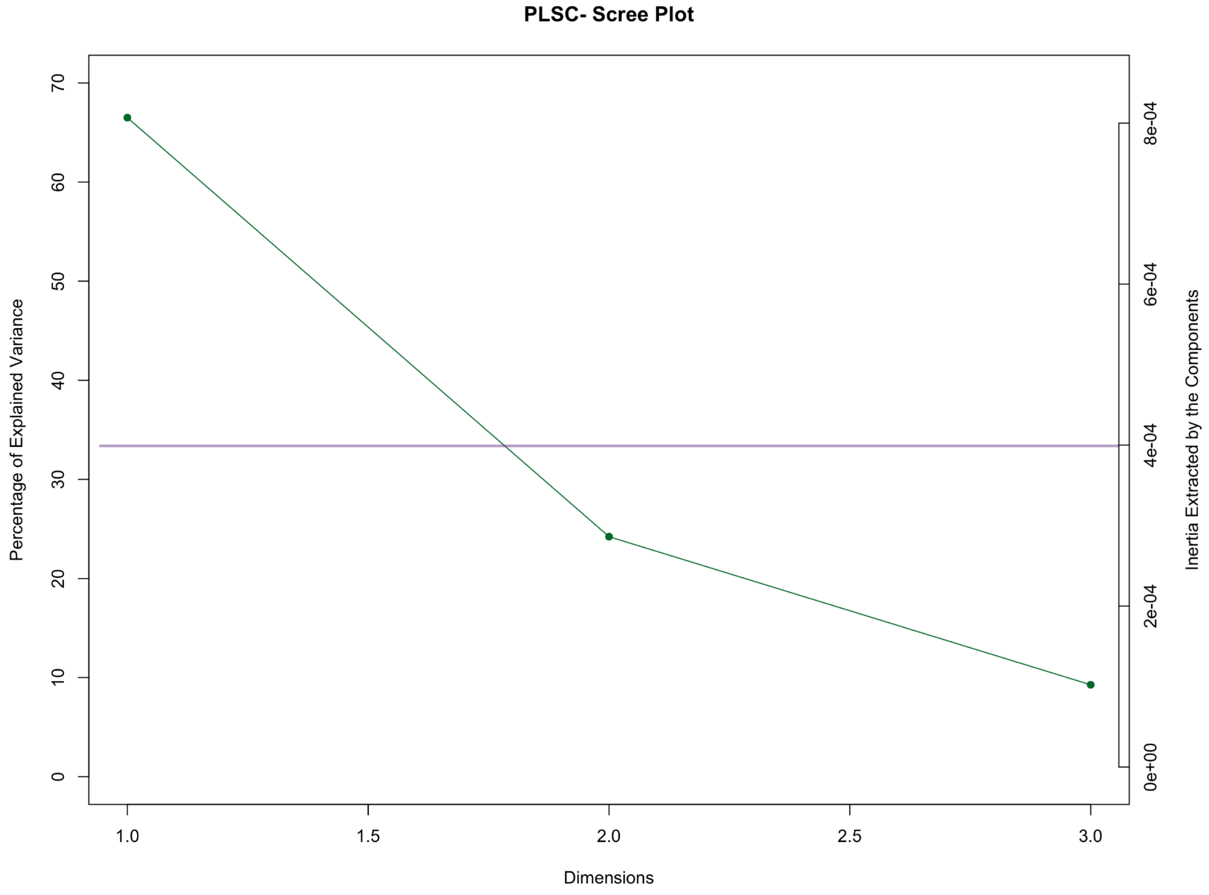

**Supplemental Figure 10.** Scree plot depicting the eigenvalues for hippocampal head, body, and tail volume and PM2.5 components analyses. The first latent dimension is identified as significant, indicating the primary sources of shared variance between the datasets. The elbow point suggests the optimal number of dimensions to retain in the model.

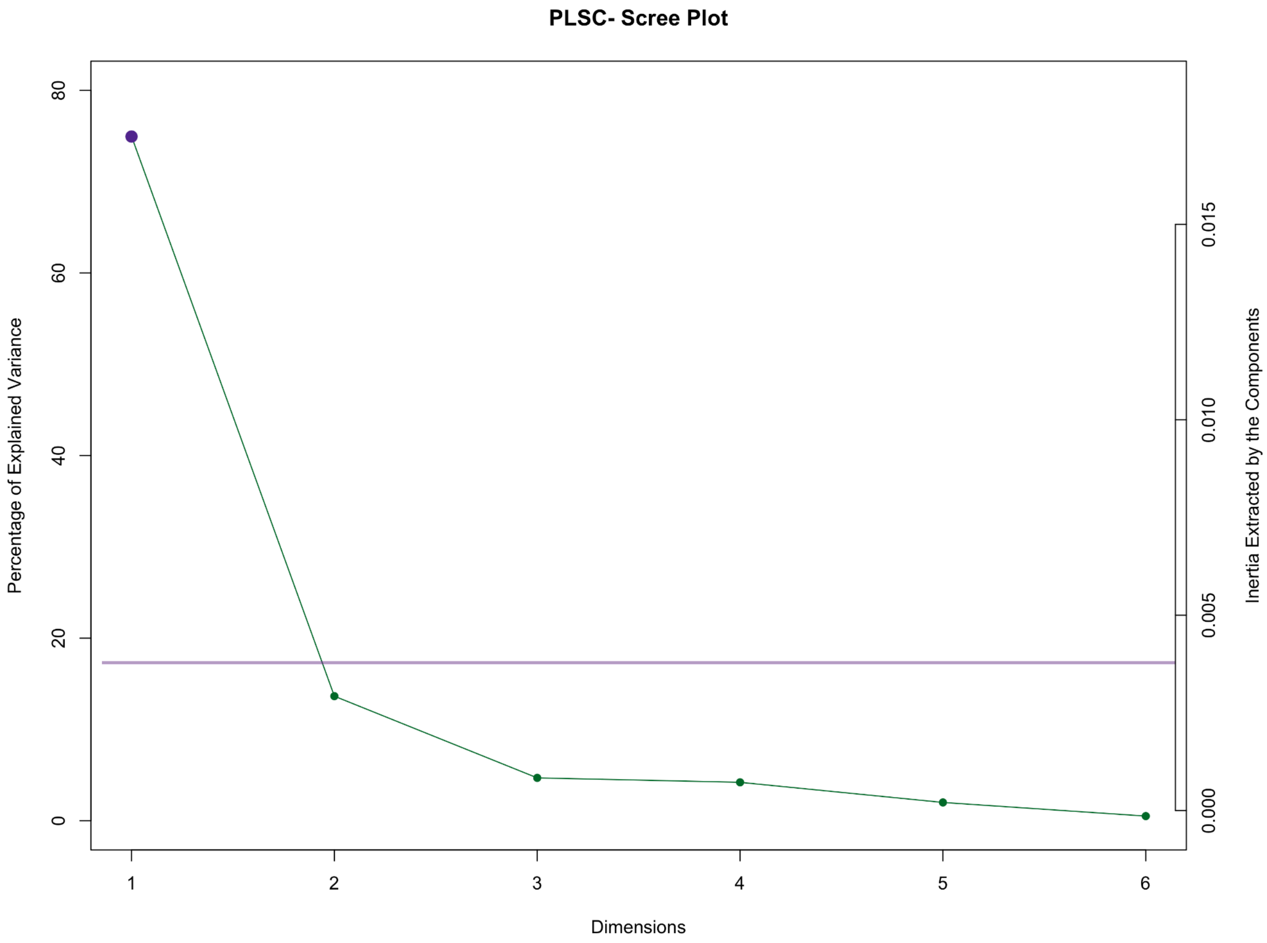

**Supplemental Figure 11.** Scree plot depicting the eigenvalues for hippocampal head, body, and tail volume and PM2.5 source analyses. The first latent dimension is identified as significant, indicating the primary sources of shared variance between the datasets. The elbow point suggests the optimal number of dimensions to retain in the model.

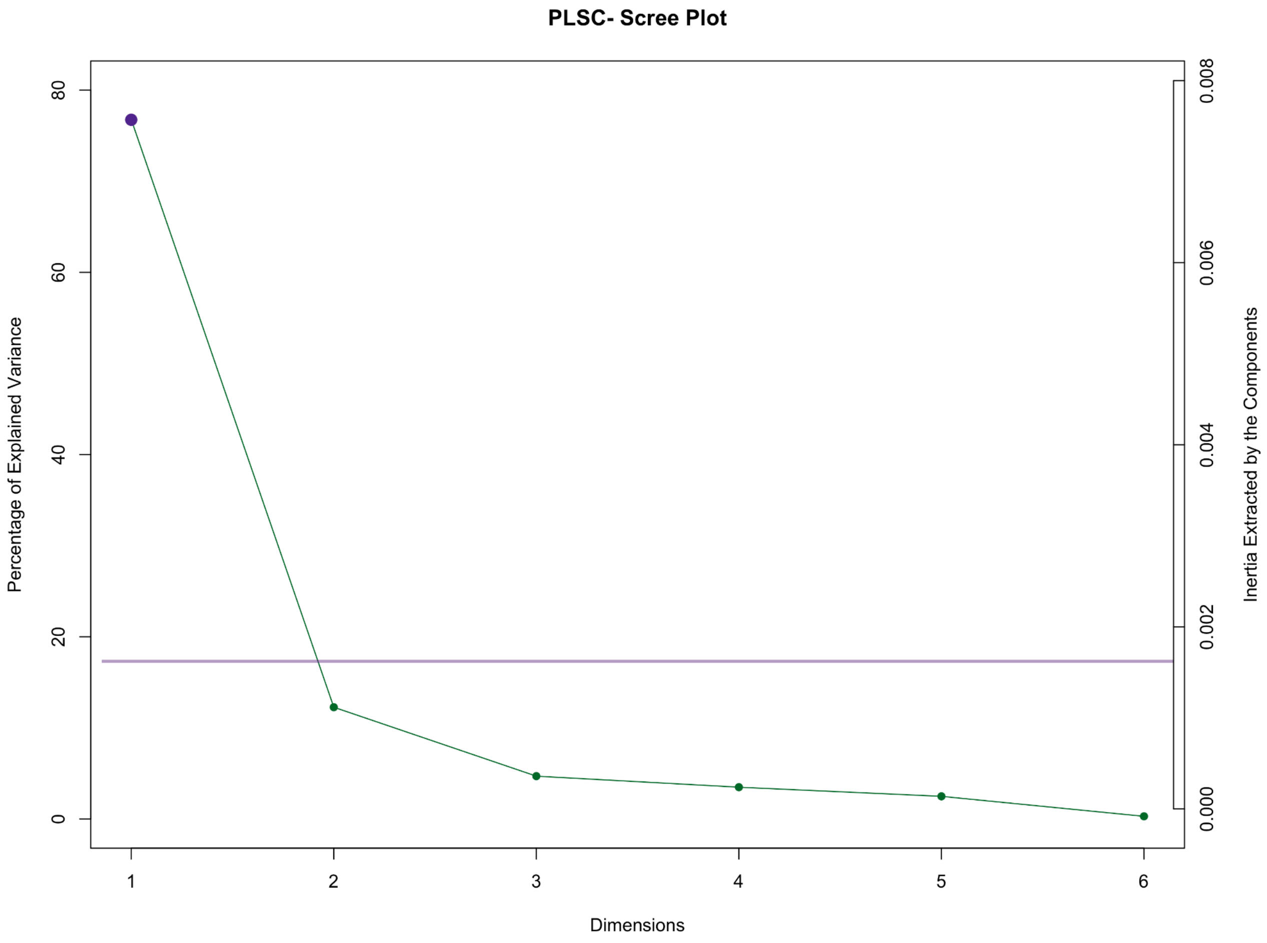

**Supplemental Figure 12.** Scree plot depicting the eigenvalues for Rey Auditory Verbal Learning Test performance and PM2.5, $NO_2$, and $O_3$ analyses.There were no significant latent dimensions.

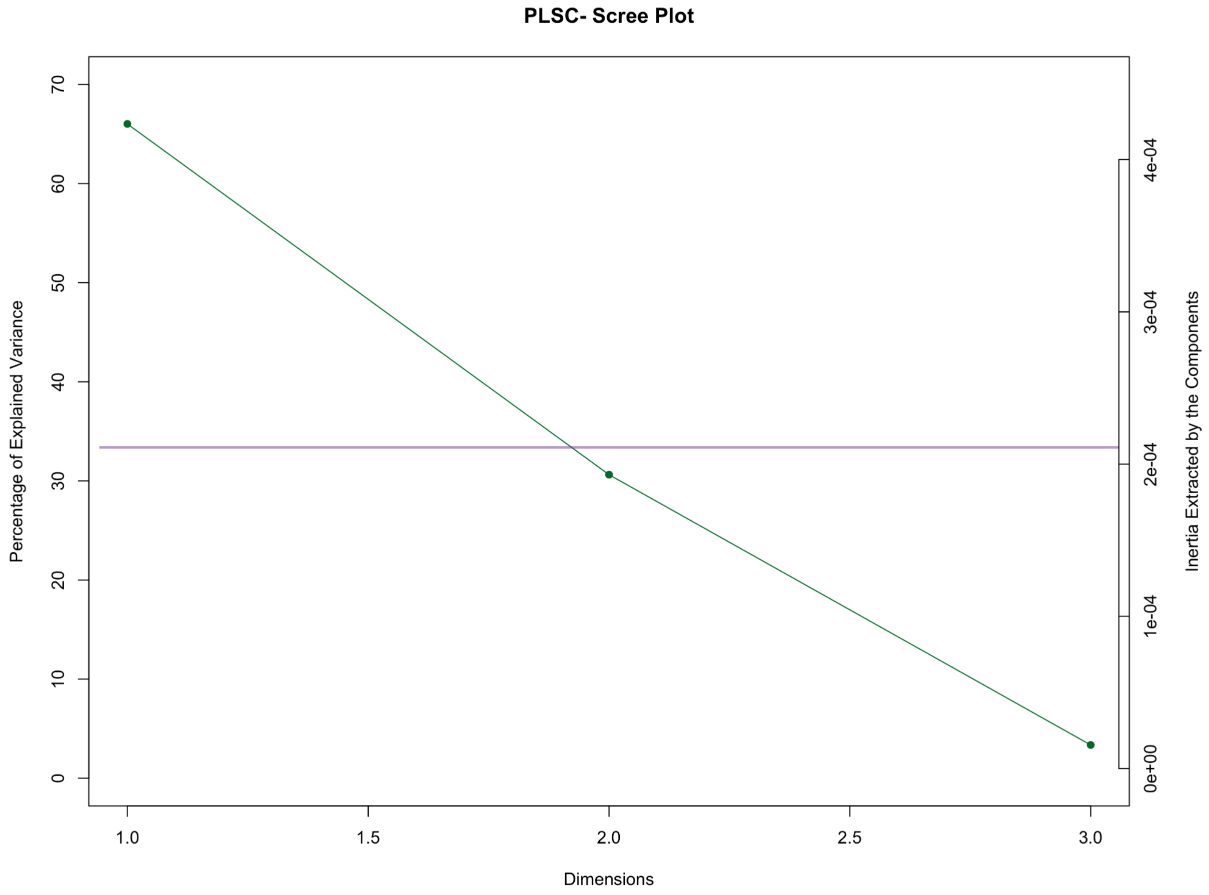

**Supplemental Figure 13.** Scree plot depicting the eigenvalues for Rey Auditory Verbal Learning Test performance and PM2.5 component analyses. The first latent dimension is identified as significant, indicating the primary sources of shared variance between the datasets. The elbow point suggests the optimal number of dimensions to retain in the model.

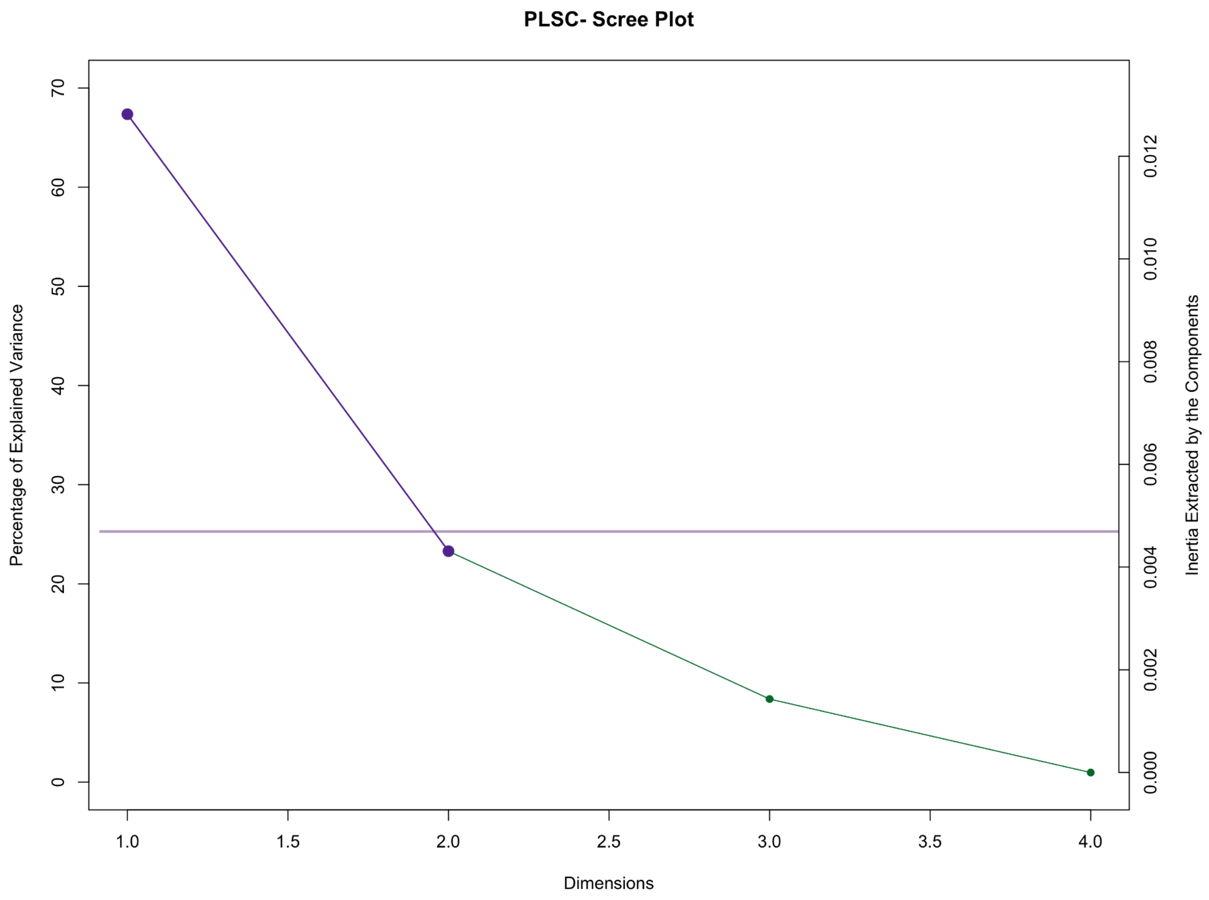

**Supplemental Figure 14.** Scree plot depicting the eigenvalues for Rey Auditory Verbal Learning Test performance and PM2.5 source analyses. The first two latent dimensions are identified as significant, capturing the primary sources of shared variance between the datasets. The elbow point suggests the optimal number of dimensions to retain in the model.

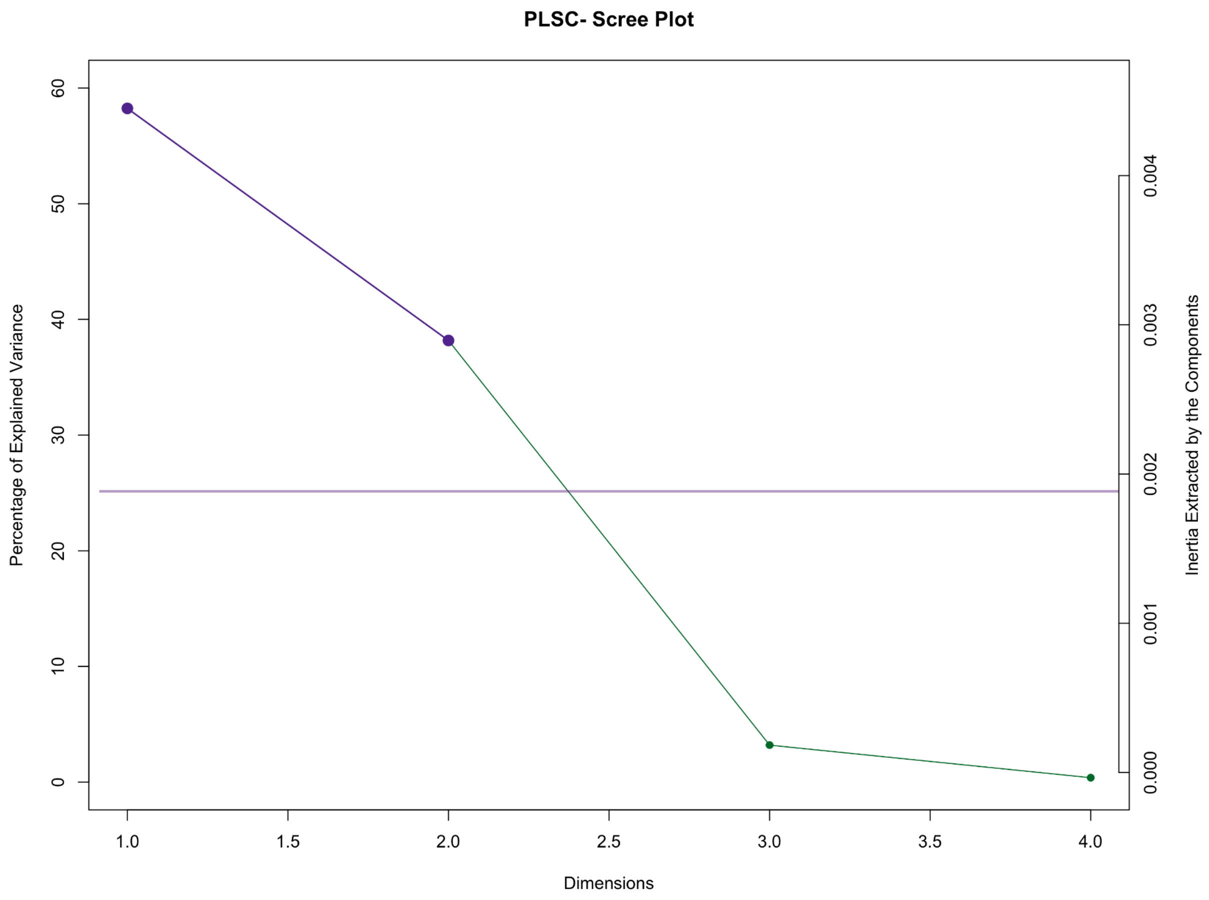

**Supplemental Figure 15. Associations between air pollution source exposure and learning and memory performance in children ages 9-11 years (n = 7,699)**. In the first latent dimension, PM2.5 sources are associated with poorer learning and immediate recall performance. In the second latent dimension, higher biomass burning is associated with RAVLT performance. RAVLT: Rey Auditory Verbal Learning Test.

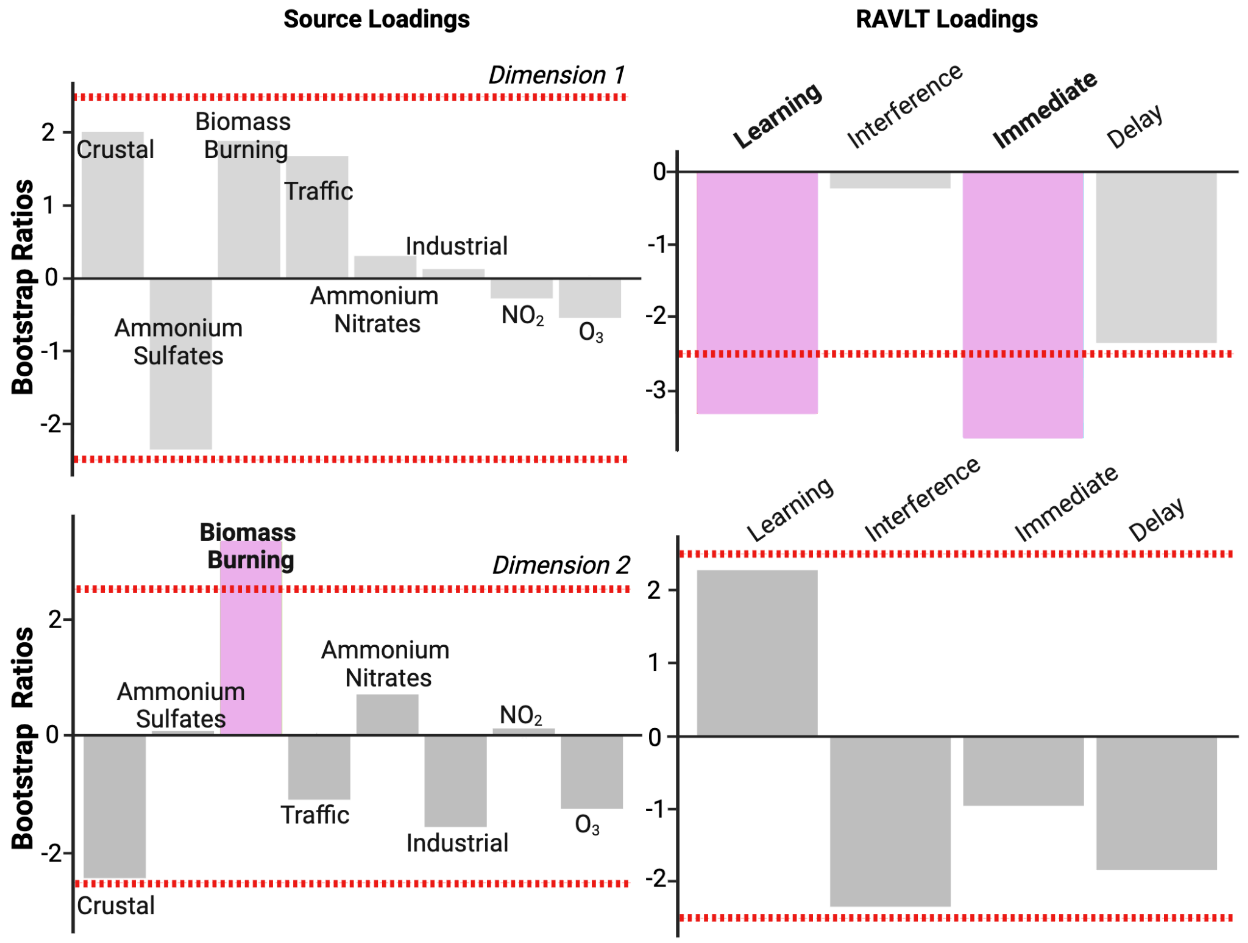

**Supplemental Figure 16.** Scree plot depicting the eigenvalues for Rey Auditory Verbal Learning Test performance and hippocampal RSI analyses. The first latent dimension is identified as significant, indicating the primary sources of shared variance between the datasets. The elbow point suggests the optimal number of dimensions to retain in the model.

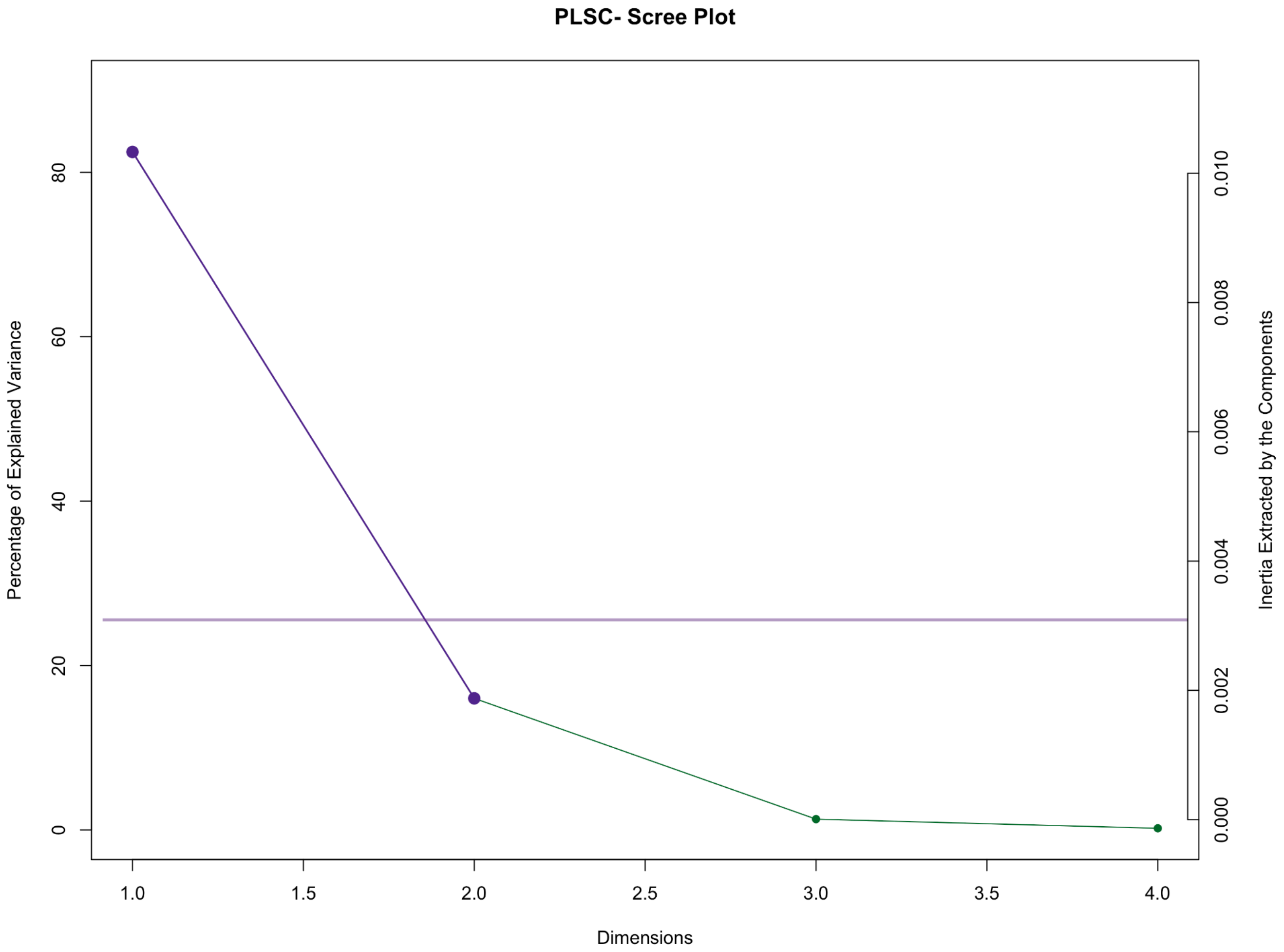

**Supplemental Figure 17.** Scree plot depicting the eigenvalues for Rey Auditory Verbal Learning Test performance and hippocampal long-axis head, body, and tail analyses. The first latent dimension is identified as significant, indicating the primary sources of shared variance between the datasets. The elbow point suggests the optimal number of dimensions to retain in the model.

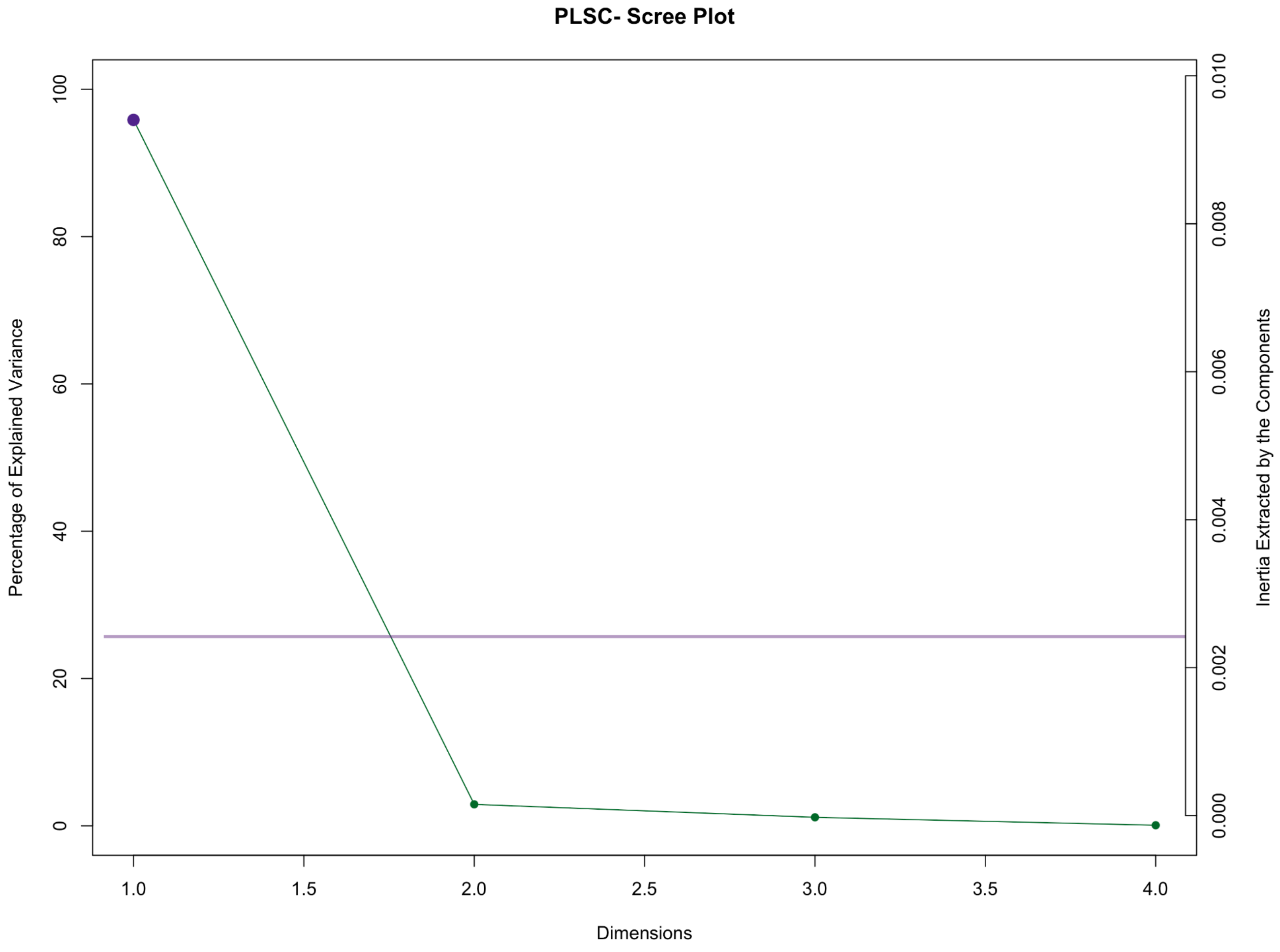
